# Theory of concentric β-barrel structures: models of amyloid beta 42 oligomers, annular protofibrils, and transmembrane channels

**DOI:** 10.1101/499061

**Authors:** Stewart R. Durell, Rakez Kayed, H. Robert Guy

**Affiliations:** Laboratory of Cell Biology, Bldg. 37 Rm 2108, National Cancer Institute, National Institutes of Health, Bethesda, MD 20892-4258, USA; UTMB Mitchell Center for Neurodegenerative Diseases, Department of Neurology, University of Texas Medical Branch, 301 University Blvd, Route 1045. Galveston, TX 77555-1045. USA; Cochiti Lake Institute for Protein Structure, 6510 Tahawash Street, Cochiti Lake, NM 87083, USA

## Abstract

Amyloid beta (Aβ) peptides are a major contributor to Alzheimer’s disease. Previously, our group proposed molecular models of Aβ42 hexamers with two concentric antiparallel β-barrels that act as seeds from which dodecamers, octadecamers, both smooth and beaded annular protofibrils, and transmembrane channels form. Since then, numerous aspects of our models have been supported by experimental findings. Here we develop a more extensive range of models to be consistent with dimensions of assemblies observed in electron microscopy images of annular protofibrils and transmembrane assemblies. These models have the following features: Dodecamers with 2-concentric β-barrels are the major components of beaded annular protofibrils (bAPFs). These beads merge to form smooth annular protofibrils (sAPFs) that have three or four concentric β-barrels. Channels form from two to nine hexamers. Antiparallel C-terminus S3 segments form an outer transmembrane β-barrel. Half of the monomers of vertically asymmetric 12mer to 36mer channels form parallel transmembrane S2 β-barrels, and S1-S2 (N-terminus and middle) segments of the other half of the monomers form aqueous domains on the cis side of the membrane. Unit cells of 42-54mers have two more transmembrane S2 segments, with four concentric β-barrels in the transmembrane region and two concentric β-barrels on the cis side of the membrane.

## Introduction

At first, second, and even third glance, the quest to determine precise molecular mechanisms by which amyloid beta (Aβ) peptides wreak havoc on the human brain appears hopeless. These peptides are shapeshifters; they assume countless forms that are often present simultaneously, and it remains unclear which of their many guises are the culprits. Much attention has focused on the most visible and stable forms: fibrils within the amyloid plaques that are the hallmark of Alzheimer’s disease (AD). Their stability has allowed the molecular structure of some forms to be determined experimentally, but even these assemblies come in multiple forms: some have U-shaped monomers ^1,2^; others have S-shaped monomers ^3–5^, some have twofold symmetry along their long axis and others have three-fold symmetry ^4,5^; and most have parallel β-sheets, but at least one highly toxic mutant has antiparallel β-sheets ^6^. However, evidence is increasing that much smaller assemblies, called oligomers, are more detrimental (reviewed in Mroczko et al., 2018 ^7^) and that those formed by the longer of the two major forms, Aβ42, are the most toxic ^8^.

A quarter of a century has passed since Arispe *et al.* ^9^ reported that Aβ peptides can form ion channels in lipid bilayers. Since then their findings have been confirmed and refined in numerous laboratories. Two recent breakthroughs support the Aβ channel hypothesis. Serra-Batiste *et al.* ^10^ have discovered conditions under which Aβ42, but not Aβ40, forms a specific β-barrel in membrane-mimicking environments. Most portions of these assemblies appear to be well-ordered and are composed of only two monomeric conformations. They find that these assemblies form highly stable ion channels with only one principal conductance (ignoring flickering currents that occurs in some, but not all, of their experiments). They have also developed a method to test the toxicity of these assemblies *in vitro* ^11^. But Aβ42 likely forms a variety of discrete channels. Bode *et al.* ^12^ found that Aβ42 oligomers, but not Aβ42 monomers, Aβ42 fibrils, or Aβ40 assemblies, form multiple types of stable nonselective channels in excised membrane patches from HEK293 cells of neuronal origin. The single channel conductance of all of these channels is greater than that reported by Serra-Batiste *et al.* ^10^. The membrane channel hypothesis for AD thus provides an explanation why antiparallel Aβ42 oligomers are more toxic and detrimental than other forms of Aβ ^13,14^. Findings that some Aβ42 oligomers have an antiparallel β secondary structure that is similar to that of OMPA (an antiparallel β-barrel channel) ^13^, that antibodies to Aβ42 oligomers also recognize some toxins that form antiparallel β-barrel transmembrane channels ^15,16^, and that the Iowa mutant that leads to early onset AD also causes some fibrils to adopt an antiparallel β structure ^6^ support our hypothesis ^17,18^ that Aβ42 oligomers, annular protofibrils, and channels possess concentric antiparallel β-barrels.

### Previous models

Several years ago we developed atomically explicit models of the structures of Aβ42 hexamers, dodecamers, an annular protofibril ^17^, and an ion channel ^18^. Many, perhaps most, large Aβ assemblies reflect their origin: *i.e*. the final structure depends upon the ‘seed’ structure from which it has grown ^16^. The starting point for our models was the hypothesis that Aβ42 hexamers can adopt a predominantly β structure with a hydrophobic core in which all monomers have well defined identical conformations and interactions with other monomers.

**Figure 1.**
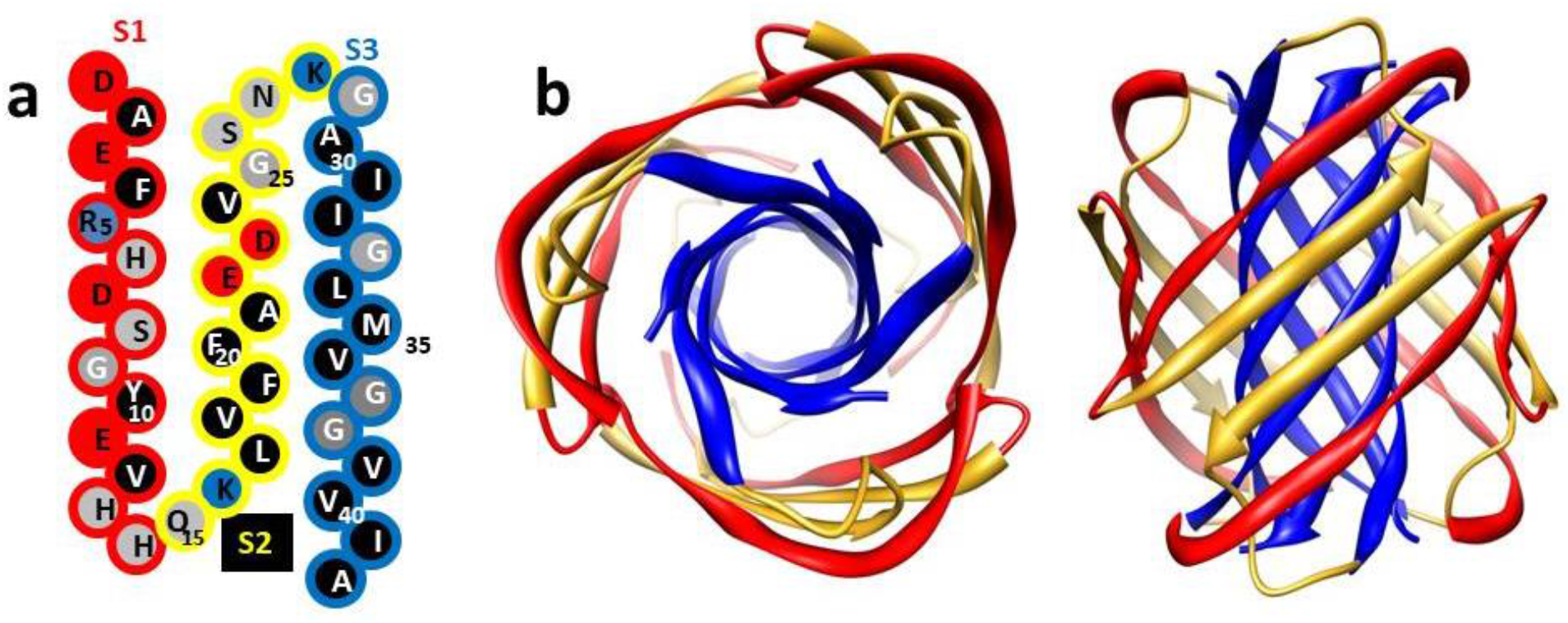
(a) Schematic representation of Aβ42 sequence. We classify the peptide into three regions of about equal length called S1 (outline color coded red), S2 (yellow), and S3 (blue). Circles are colored according to side chain polarity: red = negatively charged, blue = positively charged, grey = uncharged polar or ambivalent, black = hydrophobic. S2 and S3 segments are assigned a β secondary structure and form antiparallel β-barrels in all of our models; in most cases this pair has a U-shaped configuration similar to that observed in some fibrils ^2^. S1 forms either one or two β-strands in our models. (b) Our original model of a hexamer in which all monomers have the same conformation. The assembly has three-fold radial symmetry and 2-fold vertical symmetry. S3 segments (blue) form the hydrophobic core as an antiparallel six-stranded β-barrel. The S3 barrel is surrounded by a 12-stranded β-barrel formed by S1 (red) and S2 (yellow) segments. Copied from ^17^.

The core of our hexamer model was a six-stranded antiparallel β-barrel formed by the last third of the peptide: i.e., S3 of Fig. 1. This hydrophobic core region remained exceptionally stable in both conventional ^17,18^ and much longer coarse-grained molecular dynamic (MD) simulations ^19^. The antiparallel models were more stable than parallel models. We suggested that the first two-thirds of the peptide comprise two segments, S1 and S2, that form β-hairpins, and that these hairpins shield the S3 core by surrounding it as a 12-stranded antiparallel β-barrel (Fig. 1). As expected, these water-exposed and less compact S1 & S2 regions were less stable than the S3 core.

Several aspects of our hexamer model were unprecedented: (1) six-stranded antiparallel beta barrels had never been reported. However, Laganowsky *et al.* ^20^ have since found that a segment from an amyloid-forming protein, alphaB crystalline, indeed has the six-stranded antiparallel β-barrel motif (which they call Cylindrin) and Do *et al.* ^21^ found that several eleven-residue peptides with the sequences of portions of S3 that includes methionine also form this motif. Their calculations confirm our findings that the presence of glycine facilitates packing of aliphatic side chains (especially methionine) from adjacent monomers in the interior of the barrel. The importance of these residues is supported by findings that mutation of Gly 33 to Ala ^22^ or oxidation of Met 35 ^23^ reduces toxicity and alters oligomerization of Aβ42. Also, MD simulations of randomized clusters of small peptides with sequences of the central portion S2 of Aβ ^24^ and with the sequence of a portion of S3 ^25^ form 6-stranded β-barrels in some, but not all, simulation runs. (2) Our β structures were antiparallel whereas all known Aβ fibril β-structures were parallel. However, since then an Iowa mutant responsible for some forms of early onset AD has been shown to form fibrils with antiparallel β-sheets ^6^. More important, recent experiments indicate that some Aβ42 oligomers do have an antiparallel β secondary structure that is similar to that of OMPA ^13^ (an antiparallel β-barrel channel), and antibodies that recognize Aβ42 oligomers also recognize some toxins that form antiparallel β-barrel transmembrane channels ^15,16^. Also, antiparallel oligomers are more toxic than those with parallel structures ^13,14^. (3) Concentric β-barrels had never been observed when we proposed the structures. But recent studies have found that the channel-forming toxins Areolysin ^26^ and Lysenin ^27^ do in fact contain concentric β-barrels. (4) There was no experimental evidence supporting our proposal that the S1 segments form a β-strand or possibly a β-hairpin. Subsequently two fibril structures with S-shaped monomers that include S1 have been solved (one with 3-fold symmetry ^4,5^ and one with two-fold symmetry ^3^). In both of these, the S1 and S2 segments comprise a parallel β-sheet with a bend near the center of S1. Although often ignored by modelers, numerous findings indicate that alterations within S1 segments affect the toxicity of Aβ (see Experiment Tests section). (5) There was no experimental evidence that Aβ42 forms β-barrels. However, Serra-Batiste *et al.* ^10^ have recently discovered membrane-mimicking conditions under which Aβ42 oligomers form a well-defined β-barrel composed of only two monomeric conformations.

## Part 1: Theory and rationale for concentric β-barrel models of circular Aβ42 assemblies

Here we expand our initial models to include new schematics of concentric β-barrel models of dodecamers, octadecamers and a multitude of annular protofibrils (APFs) and channels. Models described below are based on the following: (1) β-barrel theory that allows the diameter and strand tilt angles of β-barrels to be calculated quite accurately from the number of strands, N, comprising the barrel and the sheer number, S, which is related to how much the strands tilt relative to the central axis of the barrel (see Fig. 2 and Murzin, *et al.* ^28^ for calculation of barrel diameter and Chou, *et al.* ^29^ for calculation of the tilt angle); (2) experimentally determined distances between β-sheets of Aβ fibrils and between adjacent concentric β-barrels; (3) EM images of sAPF that show the size of the assemblies, the thickness of the circular wall as viewed from the top of some images, and an apparent 6-fold (and occasionally 5-fold, 7-fold, and 8-fold) radial symmetry of some images (the last two properties have not been reported previously); (4) a preference for structures that are similar to those of fibrils; (5) the hypothesis that sAPFs develop from the oligomer structures described above and retain radially and for APFs vertically symmetric antiparallel β-barrel and multiple-of-six-monomers properties of the seed structures; (6) a preference for assemblies that are symmetric and have few monomeric conformations; and (7) a preference for the simplest explanation (Occam’s Razor).

### Concentric B-barrel theory

Strands of almost all known β-barrels are tilted and spiral around the central axis in a right-handed manner. Diameters of β-barrel backbones and tilt angles of the strands were calculated from the number of strands (N) and the sheer number (S), as illustrated in Fig. 2. The S/N ratio ranges from 1.0 to 2.0 for smaller barrels, with values as low as 0.0 for larger barrels ^30,31^.

**Figure 2.**
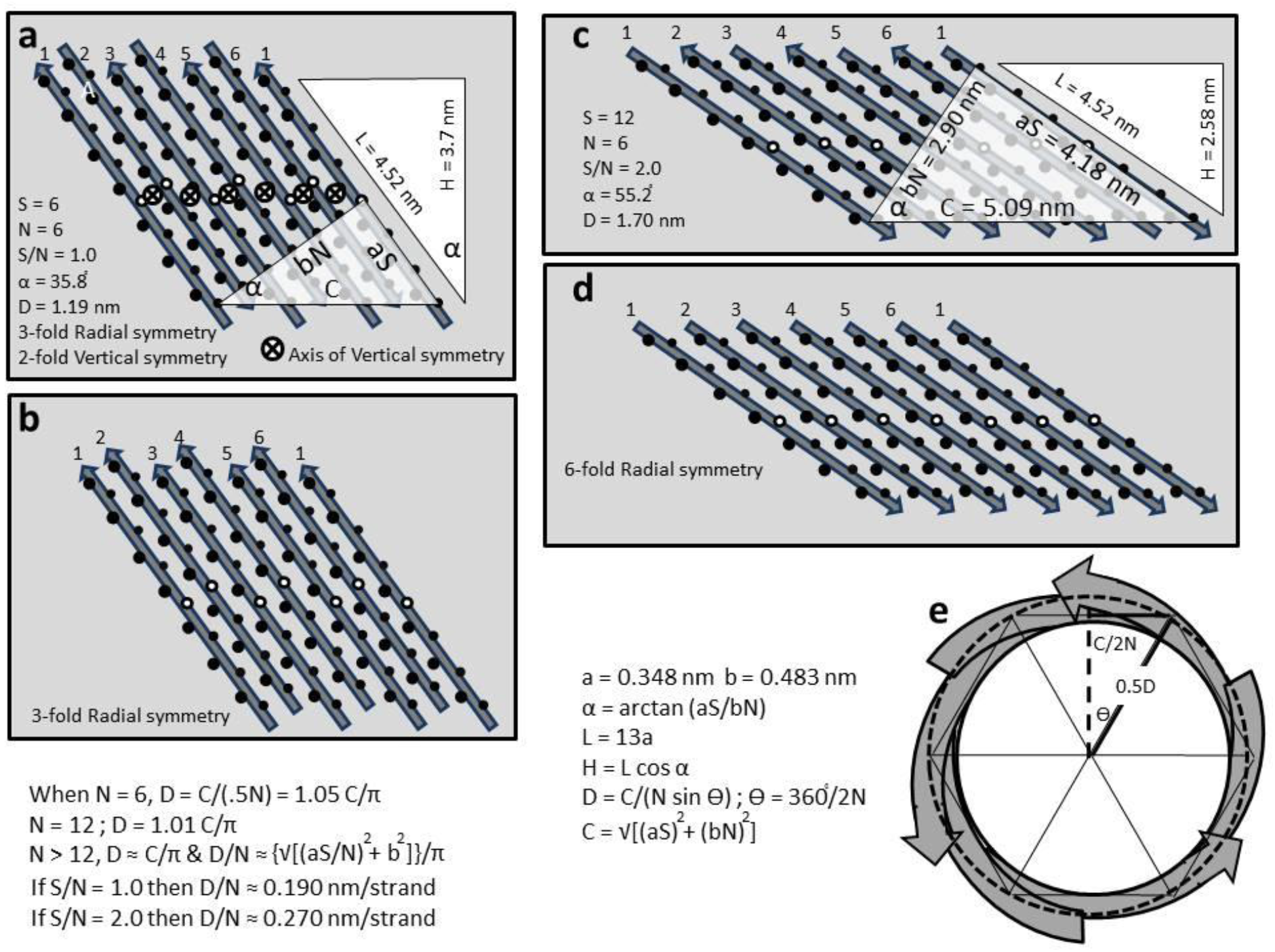
Representations of six-stranded β-barrels depicted as if the barrel were split open, spread flat, and viewed from the inside of the barrel. Circles represent side chains on the front (interior) (larger to the left) and back (exterior) side (smaller to the right); the white circles represent the strand’s midpoint. The ID numbers of the strands are shown above, with strand #1 shown twice. (a) Antiparallel barrel with a sheer number, S, of six and 3-fold radial symmetry and 2-fold vertical symmetry. (b) Parallel β-barrel for which S/N = 1.0 with 3-fold radial symmetry. (c) Antiparallel β-barrel for which S/N = 2.0 with 3-fold radial and 2-fold vertical symmetry. (d) Parallel β-barrel for which S/N = 2.0 and 6-fold radial symmetry. S can be determined by the number of residues on the right side of the shaded right triangles in (a) and (c) for the end strand. The length of this side of the triangle equals the distance between adjacent residues on the same strand (a = 0.348 nm) times S. The length of the top side of the triangle equals the perpendicular distance between strands (b = 0.483 nm) times the number of strands (N = 6). The strand tilt angle, α = arctan(aS/bN). C is the hypotenuse of the triangle and approximates the circumference of the barrel when N is 12 or greater. (e) Representation of a symmetric 6-stranded antiparallel β-barrel (same as model in (a)). The dashed line represents the circumference of the barrel. The diameter of a small barrel is calculated more accurately by D = C/[N sin(360°/2n)] as illustrated in (e).

### Symmetry

When sequentially identical subunits comprise a protein, they usually have identical conformations and interactions with neighboring subunits ^32,33^. This is true for the core region of all experimentally-determined Aβ fibril structures, for our β-barrel models of hexamers, and for experimentally-determined transmembrane channel structures. Symmetry and lattice constraints are often essential for the determination of protein crystal structures and simplify modeling of multi-subunit proteins. For cases in which each β-strand of a β-barrel belongs to a different but sequentially identical monomer, all monomers of an antiparallel β-barrel can have identical conformations and interactions only when S/N = 1 or 2, and for parallel β-barrels only when S/N = 2; otherwise some strands are staggered relative to their neighbors (Fig. 2). Most of the models we currently favor have these S/N values. For assemblies with concentric β-barrels, symmetry among the barrels is also important. Our models are constrained substantially by requiring the axes of overall symmetry to be the same for all of the barrels. Our models of oligomers and sAPFs have 2-fold vertical symmetry and a maximum of M/6-fold radial symmetry, where M is the number of monomers in the assemblies.

### Relationships among X-fold symmetry, number of strands (N), and sheer number (S)

If a β-barrel has X-fold radial symmetry formed from X unit cells that each contain Y β-strands, then N = XY where N in the number of strands within the barrel. If each radial unit cell has 2-fold vertical symmetry, then Y must be an integer multiple of two. Adjacent β-strands can shift relative to each other in steps of two residues. The number of such shifts, Z, determines the tilts of the strands and thus the sheer number, S. If radial symmetry is maintained the shifts must occur within or between each unit cell. In either case, the S/N ratio must be 2ZX/XY. For example, if the barrel has X = 6-fold radial symmetry and each radial unit cell has Y = 8 strands, then N = XY = (6)(8) = 48, S = 2XZ, and S/N = Z/4. The number of S/N values increases as the number of strands per unit cell increases. For example: if Y = 1, then S/N must be 2.0; if Y = 2 then it can be 1 or 2; if Y = 4 it can be 0.5, 1, 1.5, or 2.0; if Y = 3 then S/N can be 2/3, 4/3, or 2.0; if Y = 6 S/N values of 1/3, 1.0, 5/3 are also possible. S/N values become more restrictive for concentric β-barrel structures in which all barrels have the same symmetries, the distance between barrels is constrained, and the diameters and shapes of the assemblies are required to be consistent with EM images. In most symmetric concentric β-barrel assemblies considered here, only one set of N and S/N values is possible.

### Distance between β-barrels

We have relied on known structures to approximate an acceptable range of distances between concentric β-barrels. There are only two instances in which structures of concentric β-barrels have been determined experimentally. The best described, Aerolysin, has seven-fold radial symmetry ^26^. Its inner 14-stranded antiparallel β-barrel has 7 unit cells with two strands per cell and an S/N value of 1.0. It is surrounded by a 21-stranded β-barrel for which S/N = 4/3 and each of the 7 unit cells has three antiparallel strands. The backbones of the two barrels are about 0.9 nm apart. For Lysenin ^27^ the gap distance is ~1.2 nm between an 18-stranded antiparallel β-barrel (X = 9, Y=2, S/N = 1.0) and a 27-strand β-barrel (X = 9, Y = 3, S/N = 4/3). The distance between β-sheets of Aβ fibrils is about 1.0 nm ^2^.

The diameter of barrels for which S/N = 1.0 can be approximated as 0.19N nm (Fig. 2). If N increases by 12 for each successive barrel, then the gap distance between barrels will be (12)(0.19)/2 = 1.14 nm. This is at the upper end of the expected value, but a slightly wider gap may leave some space for hexane (used to develop APFs) in APFs and lipid in channels. The presence of hexane could facilitate packing arrangements between barrels that change as the size of the APF changes, and may be left over when hexane-filled dodecamers merge.

### Effects of Seeds

Most, if not all, ordered Aβ assemblies are influenced by “seed” structures from which they develop. The putative seed structure for the models presented here is a hexamer with two concentric β-barrels that have two-fold vertical symmetry and three-fold radial symmetry for both barrels. We propose that dodecamers and sAPFs retain or expand these symmetries, that channels retain the radial symmetries, and that these assemblies all have integer multiple of six monomers.

### Assumptions about Aβ42 β-barrel structures

The S3 barrels are the simplest to model; they typically have only one monomer conformation within the barrel, an integer multiple of six strands, and 2-fold vertical symmetry. The preceding two thirds of the peptide pose the greatest challenge. We divide this region into three segments (S1a, S1b, and S2) that always have a β-strand structure. S2 strands are always components of a β-barrel in our models, and S1a and S1b often are (Fig. 4). These segments are connected by residues [(D-S-G) connecting S1a to S1b, (H-H-Q-K) connecting S1b to S2, and (D-V-G-S-N-K-G) connecting S2 to S3; D, S, G, and N occur frequently in turns and random coils] that can adopt a variety of conformations, allowing formation of the types of β-strands, β-hairpins, and β-U-turns that will appear in our models (Fig. 3).

**Figure 3.**
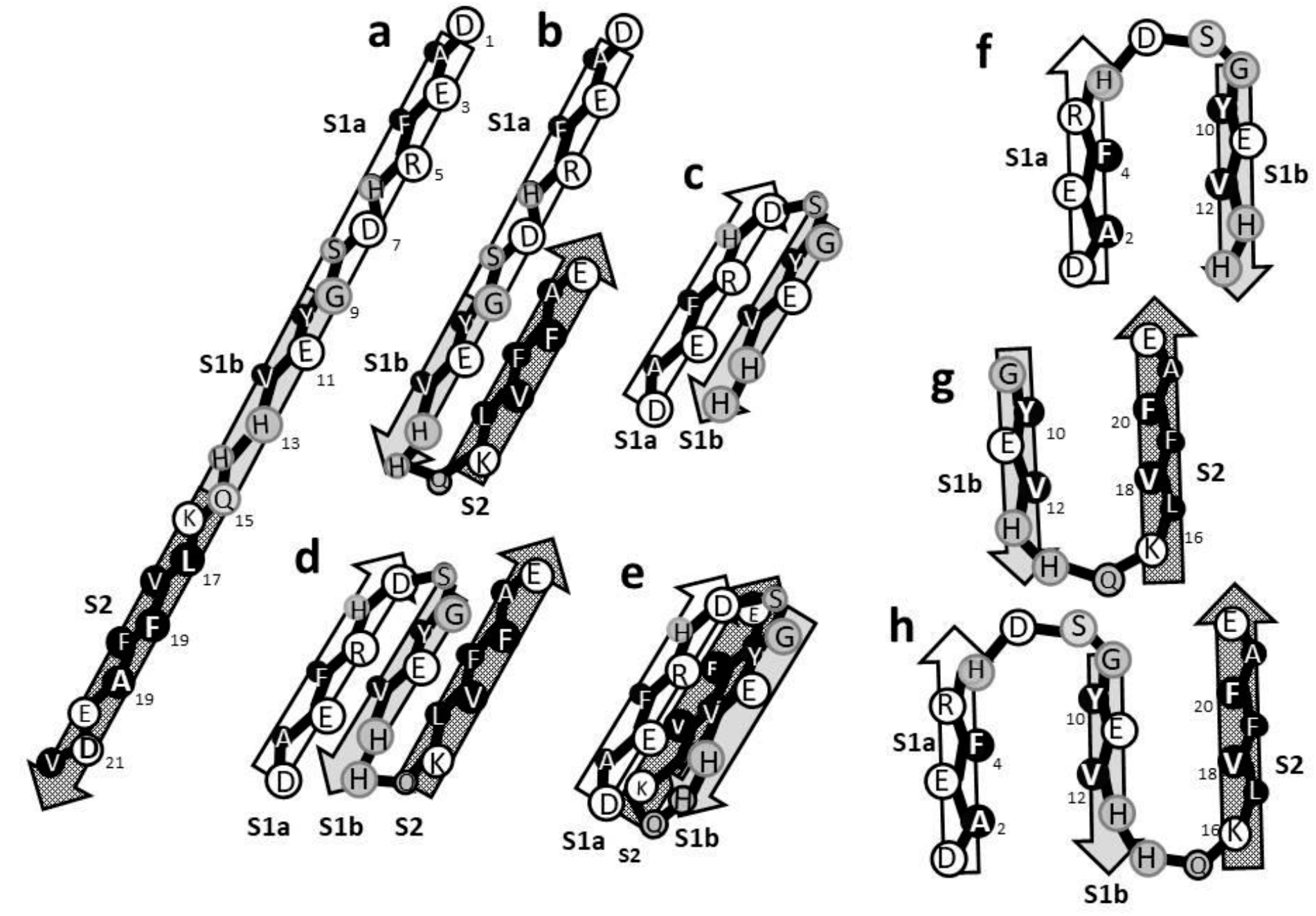
Representations of β folding motifs for the initial portion of the Aβ peptide used in our models. The strands are tilted as they would be in a β-barrel. The larger circles on the right side would be oriented toward the viewer. (a) A continuous β-strand. (b) A β-hairpin formed by S1b and S2. (c) A β-hairpin formed by S1a and S1b. All hydrophobic side chains (black) are on one side of the hairpin and all charged side chains (white) are on the opposite side. (d) A 3-stranded structure formed by S1a-S1b-S2. (e) A S1a-S1b β-hairpin over a S2 β-strand (smaller letters). (f-h) U-turn structures in which the strands [(f) S1a-S1b, (g) S1b-S2, and (h) S1a-S1b-S2] are each in a different β-barrel. Note that hydrophobic side-chains in black are buried between the strands with the exception of a putative salt-bridge formed between E11 and K16 of the double U-turn structure of 3h, which would be a triple U-turn structure were S3 included.

S2 and sometimes S1a and S1b comprise the aqueous-exposed β-barrels in our modes. Fig. 4 illustrates side views of portions (radial unit cells) of these barrels spread flat; the structures of a, b, and c have S/N values of 1.0; S/N = 2.0 for the structure in d. Fig. 4a shows the arrangement we now propose for the hexamer, where the S1a-S1b-S2 strands form a 3-stranded β-sheet and adjacent sheets are antiparallel. The outer barrel of one dodecamer model and some of the smaller sAPF models described below do not include S1a (Fig. 5b).

Fig. 4c illustrates the radial unit cell of the outer barrel of models that exhibit radial symmetry. The central region (semi-transparent parallelogram) of the unit cell is formed by antiparallel S2 strands while the sides contain S1a and S1b strands. The S2 strands form S1b-S2 β-U-turns with two hydrophobic side chains of S1b strands (lower portion with parallelogram) interacting with two hydrophobic side-chains in the center of S2. These S1b layers do not form β-barrels, but produce radially symmetric protrusions on the exteriors of some sAPFs.

We advocate that sAPFs with no radial symmetry have outer β-barrels composed exclusively of antiparallel S2 strands that may be flanked on each end by S1a-S1b β-hairpins (Fig. 4d).

**Figure 4.**
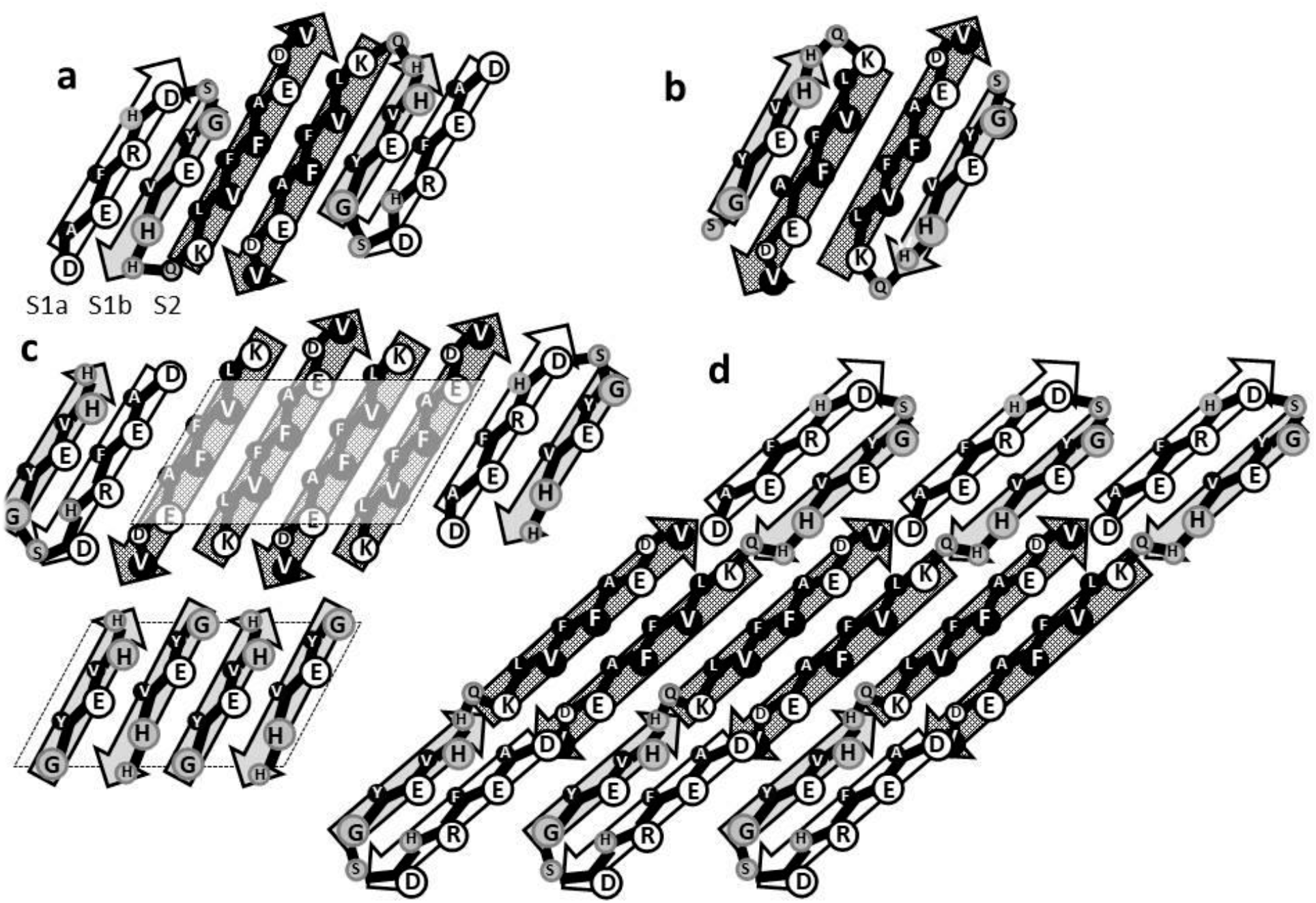
Illustration of the β-strands of outer barrel unit cells viewed from the exterior and spread flat. Larger circles on the right side of the strands represent exposed side chains. All cells have 2-fold vertical symmetry and S/N values of 1.0 (a, b, & c) or 2.0 (d). (a) The arrangement proposed for soluble hexamers in which each monomer contributes three strands to the outer barrel. (b) The arrangement proposed for the dodecamer of Fig. 5c and some small sAPFs with each monomer contributing S1b and S2 β-strands. (e) Model for the outer surface of sAPF that exhibit radially symmetric irregularities on the perimeter. Antiparallel S2 segments of the unit cell are flanked by S1a and S1b segments. The S2 strands are part of S1b-S2 β-hairpins and the antiparallel S1 β-sheet at the bottom protrudes from each unit cell. (d) The arrangement proposed for some larger sAPFs where the central part of the outer barrel is formed exclusively by S3 strands. S1a-S1b β-hairpins flank the S2 β-barrel.

## Part 2: β-barrel models of Aβ42 oligomers

Crosslinking and other methods indicate that primary oligomers of Aβ42 are hexamers, or multiples of hexamers: dodecamers (12mers) and octadecamers (18mers) ^23^. The schematics of Fig. 5 illustrate how hexameric β-barrel structures could interact to form dodecamers with differing sizes and shapes. In our initial models, we postulated that S1 forms a single β-strand. However, we now favor antiparallel β-barrel models that have S/N values of 1.0 and that have twelve more strands than the barrel it surrounds. If this is the case for hexamers, then the 6-stranded S3 core β-barrel should be surrounded by an 18-stranded β-barrel formed by the other segments. Figs. 4a and 5a illustrate how the S1a-S1b-S2 strands could form a 3-stranded β-sheet and that six such sheets could form an 18-stranded β-barrel that has an interior lined with hydrophobic side chains (A2 and F4 of S1a, Y10 and V12 of S1b, and L19, F21, and A23 of S3) and a hydrophilic exterior (except for V20 and F22 of S2).

Figs 4b illustrates that triple stranded S1a-S1b-S2 sheets could form a β-barrel large enough to surround the hydrophobic core of a dodecamer comprised of three concentric β-barrels. Previously we developed atomically explicit versions of this model ^17^, except for the outer barrel which had continuous S1 strands instead of S1a-S1b β-hairpins. The six-stranded antiparallel S3 core of the dodecamer was highly stable during MD simulations, the 12-stranded middle layer S2-S3 barrel was slightly less stable, and the outer S1-S2 layer was the least stable.

**Figure 5.**
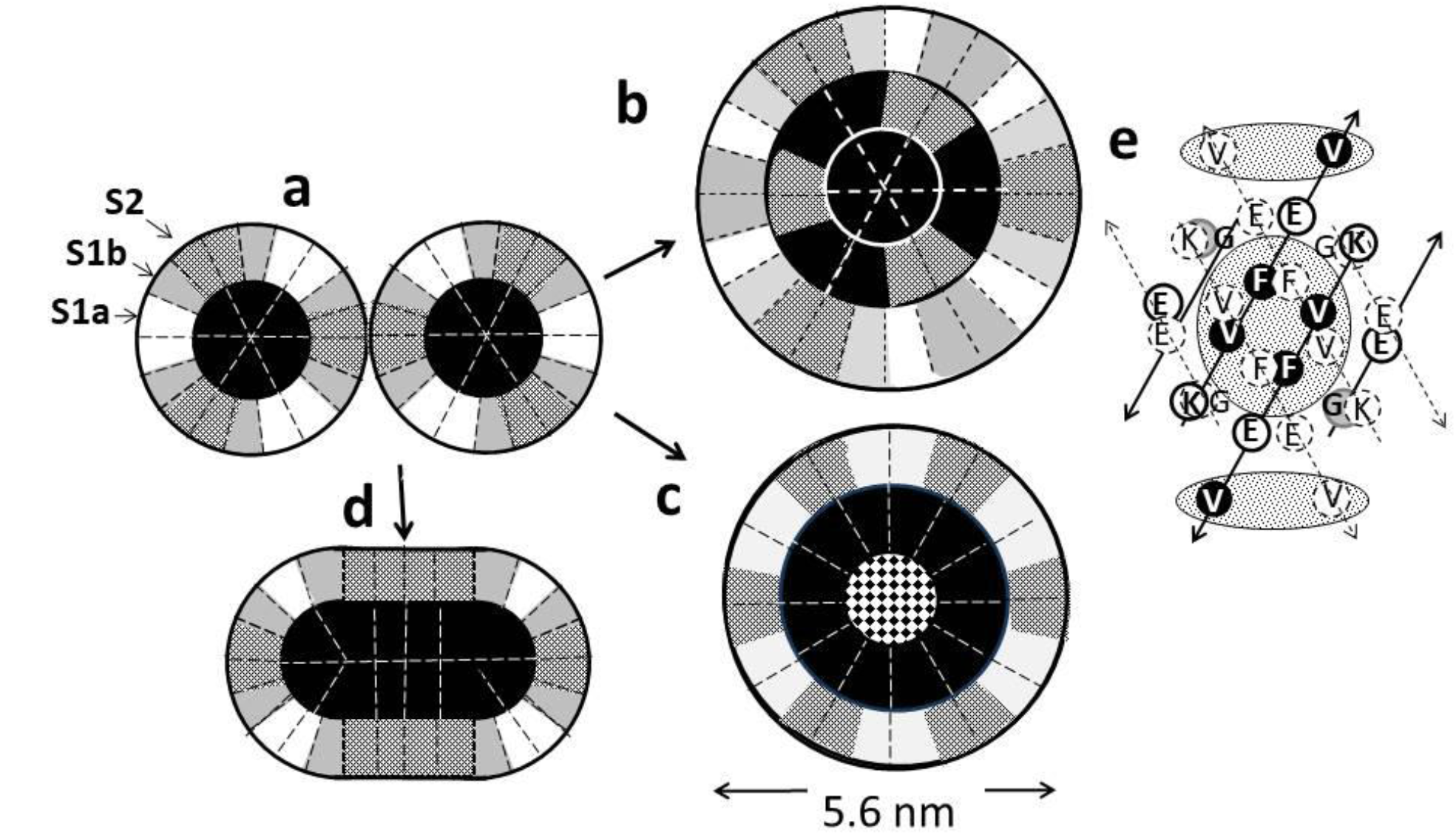
Illustrations of how Aβ42 hexamers may interact to form dodecamers. Each circle represents an antiparallel β-barrel with 2-fold vertical symmetry and S/N = 1.0 as viewed from the top. Dashed lines separate monomers. (a) Models of two hexamers interacting. The black circle in the center represents a hydrophobic 6-stranded antiparallel S3 β-barrel. This barrel is surrounded by an 18-stranded antiparallel barrel formed by S1a (white), S1b (gray) and S2 (stippled) strands. (b) Model of a dodecamer that forms in the absence of hydrophobic molecules (*e.g*., hexane, fatty acid, or lipid). This model has three concentric antiparallel β-barrels: a core 6-stranded S3 barrel, a middle 12-stranded barrel formed by six S2 and six S3 strands, and an outer 24-stranded β-barrel formed by six S1a-S1b β-hairpins (white and dark gray covering S2 of the middle barrel, see Fig. 3e), six continuous S1 (light gray), and six S2 strands (stippled). (c) Model of dodecamers formed in the presence of hydrophobic molecules. The checked pattern in the center of the dodecamer represents hexane or some other hydrophobe. The inner barrel has twelve S3 strands and the outer barrel has twelve S1 (or S1b) and twelve S2 strands. The outer diameter shown in nanometers below the model is 1.0 nm greater than the diameter to the backbone of the outer barrel. (d) This dodecamer is similar to a flattened β-barrel in which two S3 sheets pack back-to-back (similar to the case in antiparallel Aβ fibrils) in the linear region while each end retains a curved conformation similar to half of a hexamer. This type of structure may occur during the transition from beaded APFs to smooth APFs. (e) Illustration of proposed interactions between two hexamers or two dodecamers at the axes of 2-fold symmetry between S2 segments. The hexamer on the far side is shown in bold letters with solid arrows and shaded side-chains; the hexamer on the near side has unshaded side-chains and dashed arrows. The shaded ovals in the background indicate clusters of hydrophobic side-chains.

The dodecamer model of Fig. 5c (a 12-stranded S3 β-barrel filled with hexane that is surrounded by a 24-stranded S1-S2 β-barrel) illustrates a new concept: that hexane, which is present in studies of APFs, or fatty acids may interact with S3 segments to stabilize dodecamers that have only two concentric β-barrels. This type of structure may correspond to the dodecamer seeds of “large fatty acid-derived oligomers” ^34^.

## Part 3: Annular Protofibrils

Two types of annular protofibrils have been reported: beaded APFs (bAPFs) resemble necklaces formed by a string of beads and smooth APFs (sAPFs) are more like smooth rings ^35^ (Fig. 6). These APFs were developed in the presence of hexane, with bAPFs forming initially, then gradually transforming into sAPFs. The most obvious components of the bAPFs are beads with a diameter of about six nanometers. We suspect that these beads correspond to dodecamers illustrated in Fig. 5c. Smaller beads that may correspond to hexamers and elongated thin structures (such as the flattened dodecamer of Fig. 5d) that may reflect initiation of sAPFs occur in smaller numbers, but we will concentrate on the circular dodecamers here.

**Figure 6.**
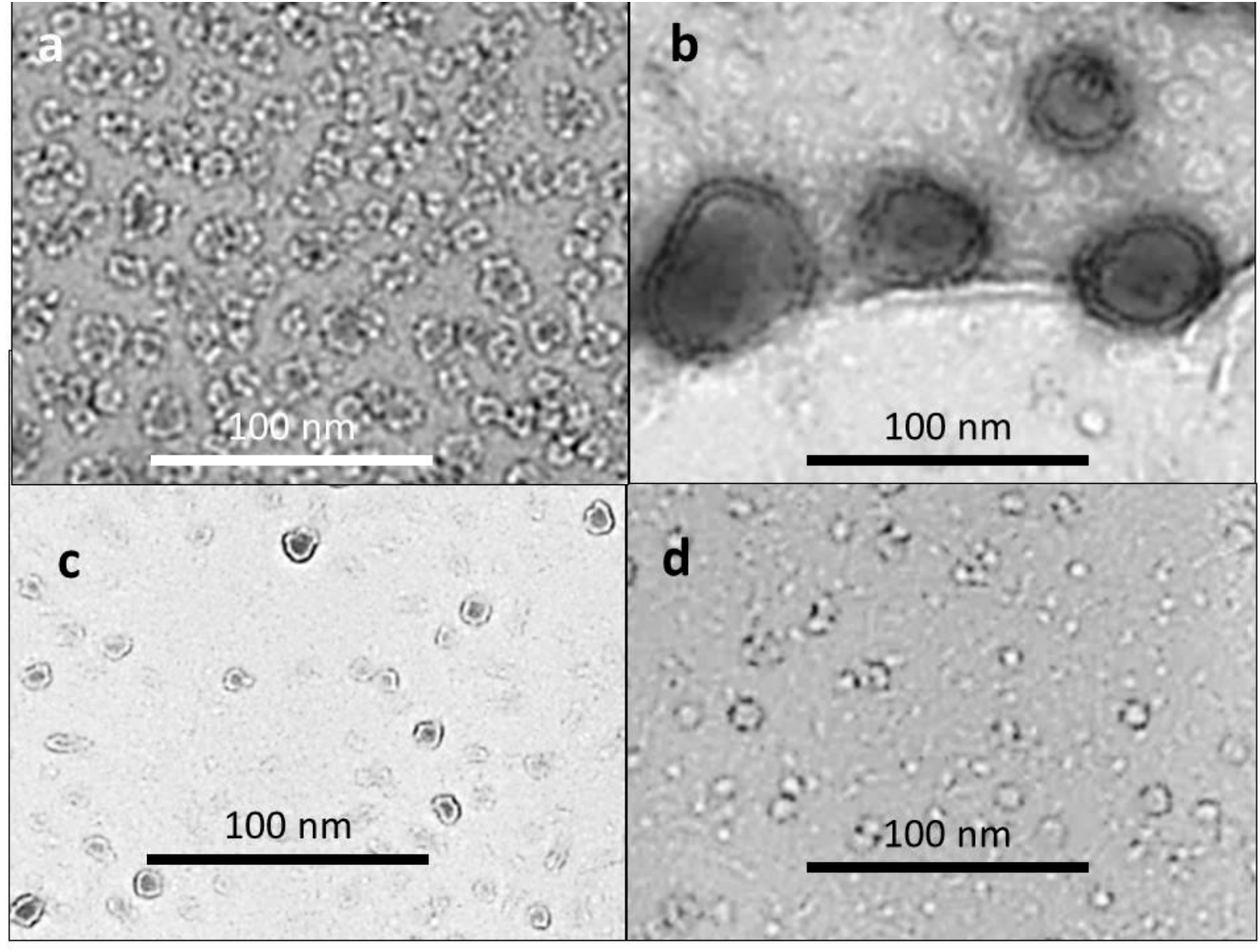
EM images of beaded (a) and smooth (b, c, & d) APFs (originals provided by Rakez Kayed). See (Kayed *et al.* ^35^) for methods.

The APFs can be classified according to their size, wall thickness, and shape (Fig. 7).

**Figure 7.**
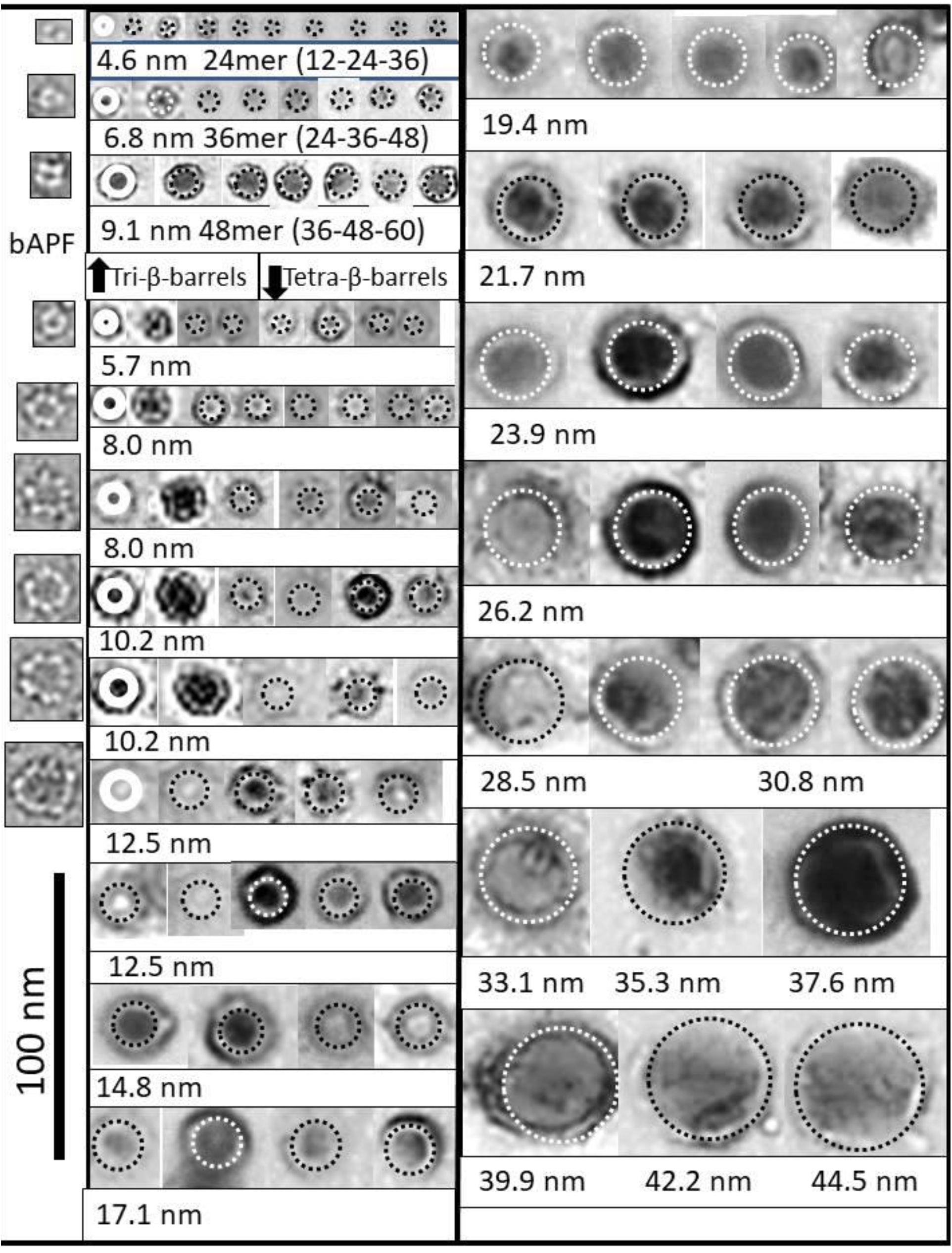
Images of smooth APFs (original EM images provided by Kayed) classified by their diameter and wall thickness and compared to beaded APFs and predictions of the diameter to the center of the wall of tri-β-barrel (first three rows) and tetra-β-barrel models. Images of putative beaded APF precursors are shown on the left for some smaller APF categories. The white solid rings illustrate the space predicted to be occupied by the β-barrel backbones drawn to scale and superimposed on the image to its right; the dashed circles indicate the predicted center of the wall if S/N = 1.0 and the number of monomers, M, is a multiple of twelve (D_wall_ = 0.19M for tri-β-barrels, D_wall_ = 0.19M/2 for Type A tetra-β-barrels in which B2 and B3 have only S3 segments, and D_wall_ = 0.19(M + 6)/2 for Type B models that have 12 S2 segments in B3 and 12 S1a-S1b β-hairpins in B4). Note that the walls of putative tri-β-barrel assemblies are thinner than those of putative tetra-β-barrel structures, and the diameters per monomer are higher. The text below the images lists the predicted value of D_wall_, which increases by ~2.3 nm for each row of tetra-β-barrels. Circles were drawn and superimposed on images using Microsoft PowerPoint. Scale bars for the EM figures were used to determine sizes of the circles and images. Additional images are shown in Fig. S1 of the Supplement.

### Beaded APF to Smooth APF

EM images of sAPFs reveal a variety of sizes and shapes. We favor three categories of concentric β-barrel models for sAPFs, as explained below. The diameter of a bAPF with X beads is only slightly greater than our models of sAPFs that have 12X monomers (Figs. 7 and 8). This relationship, plus the fact that each bead has about the same diameter as the dodecamer model of Fig. 5c, suggests that twelve monomers comprise a bead. Also, a few sAPFs appear to be in a transition state in which the beads are still visible (see images near the beginning of rows 4 to 11 in Fig. 7). Thus, we propose that beaded APFs gradually convert to smooth APFs when the beads merge to produce three or four concentric β-barrels. Some sAPFs have a two lobe peanut-shape (not shown), which suggests that sAPFs can merge to create larger sAPFs.

#### Tri-β-barrel models of sAPFs

The first category, tri-barrel models, has three concentric barrels: the central S3 barrel (B2) is sandwiched between and inner barrel (B1) and an outer barrel (B3), both formed by S1 and S2 β strands (24mer and 36mer of Fig. 8 and Fig. 9a and 9b). If S/N = 1 for each barrel, then for a 24mer, 6 S1 plus 6 S2 strands comprise B1, 24 S3 strands comprise B2, and 18 S1 plus 18 strands comprise B3, and the distance between barrels is ~1.14 nm. The diameter to the center of the wall (*i.e*., to the center of B2) is D_wall_ = (24)(0.19 nm) = 4.6 nm. Likewise, the number of strands for 36mer (Fig. 8b) and 48mer tri-β-barrels would be: 24-36-48 with D_wall_ = 6.8 nm and 36-48-60 with D_wall_ = 9.1 nm. Thus, the tri-β-barrel models are consistent with sAPFs with D_wall_ ranging from ~ 4 - 10 nm and with walls that are about 3.3 nm thick. Previously we developed an atomically explicit version of the 36mer tri-β-barrel model of Fig. 8b ^17^. The 36-stranded antiparallel S3 barrel was exceptionally stable during molecular dynamics simulations, and the backbone hydrogen bonds remained intact. The exposed inner and outer S1-S2 barrels were reasonably stable. These 163 kDa Aβ42 36mer models are candidates for the experimentally observed 150 kDa Aβ42 oligomers that form antiparallel β-sheet structures within the C-terminal S3 region ^36^.

### Tetra-β-barrel models

Our tetra-β-barrel models (propose here for the first time) for larger sAPFs with thicker walls have four concentric barrels, with two hydrophobic S3, or predominantly S3, barrels (B2 and B3) sandwiched between two hydrophilic barrels (B1 and B4) formed primarily by S2 strands (Fig. 9c-g), similar to the two S3 β-sheets that are sandwiched between two S2 β-sheets in some fibrils ^1,2^. Some intermediate-sized sAPFs display radial symmetry (6-fold 72mer and 96mer images and models of Fig. 8); whereas others appear as smooth circles (smooth images at the bottom of Fig. 8). We have developed categories of models for both types. In Type A models, B1 and B2 have the same number of strands; likewise for B3 and B4. In order for the diameter of B2 and B4 to be greater than those of B1 and B3 respectively, S/N of B2 must be greater than that of B1 and S/N of B4 must be greater than S/N of B3. Even with constraints on symmetry, distance between barrels, and the size of the EM images, there are too many adjustable parameters to be confident of a precise model. Our Type A models of smooth APFs were developed by assuming that the outer barrels B3 and B4 have twelve more monomers than the inner barrels: *i.e*., for a 60mer formed from five dodecamers 24 monomers comprise B1 and B2 and 36 monomers comprise B3 and B4. If S/N values are 72/36} 48/36} 36/24} 12/24} for the four barrels from largest to smallest, then the barrel diameters will be 9.7} 7.7} 5.5} 3.9} nm and the average distance between the barrels will be 0.97 nm. (The} symbol is used here to designate the relative positions of the concentric barrels). S values that are a multiple of 12 permit the assembly to have 6-fold radial symmetry and 2-fold vertical symmetry. The S/N values extend from 0.33 for B1 to 2.0 for B4, which are the limits of values that we have used. The S/N value of 2.0 for B4 is consistent with the illustration of the outer barrel radial unit cell in Fig. 4d.

**Figure 8.**
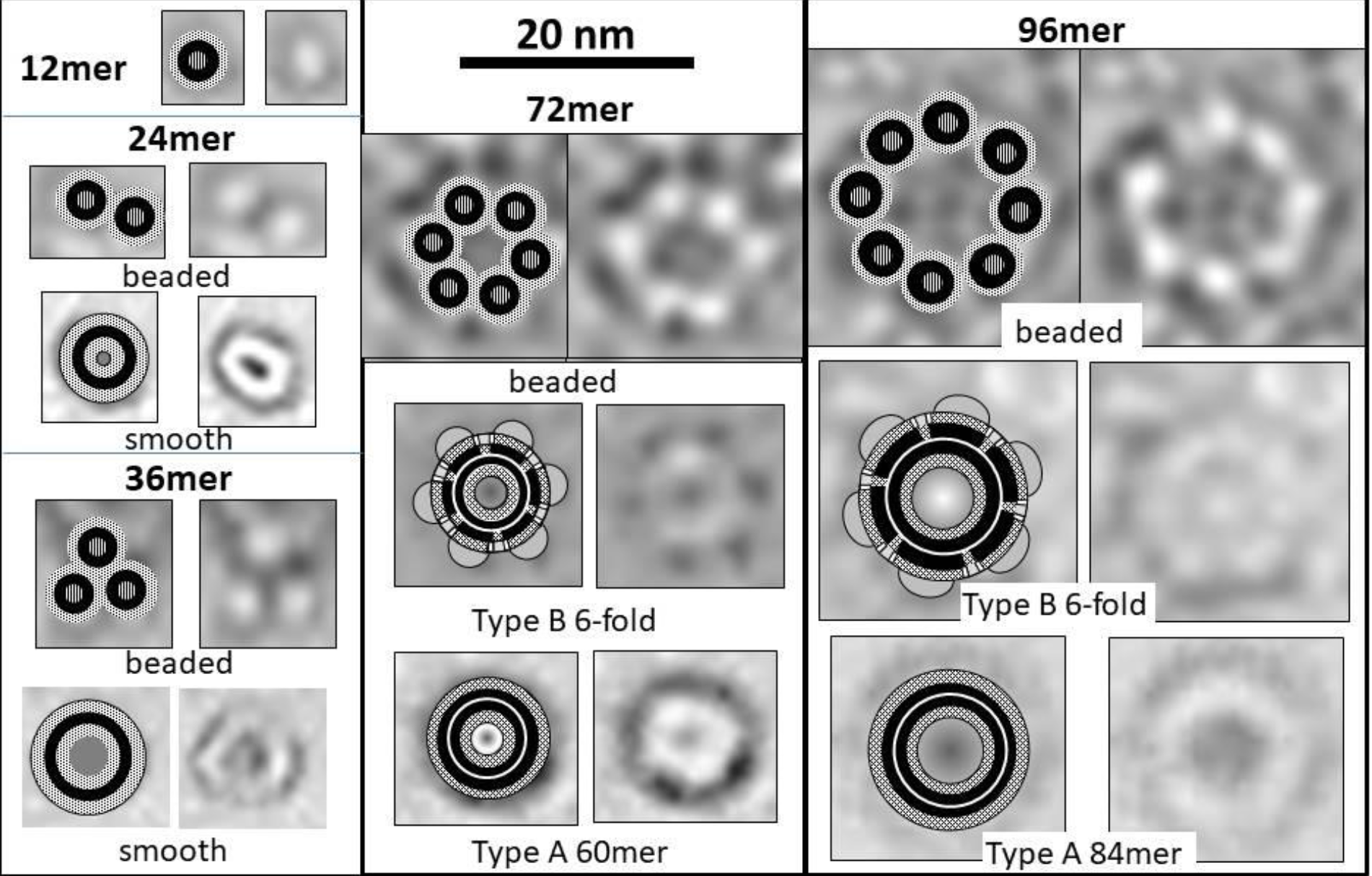
Model APFs (on left) formed by multiples of 12 Aβ42 monomers compared to images of APFs (behind models and on right). The 24 and 36mer schematics represent tri-β-barrel models with one S3 β-barrel (black) sandwiched between two S1-S2 (light-stippled) barrels. The 72 and 108mer schematics represent tetra-β-barrels with two concentric S3 β-barrels (black) sandwiched between S2 β-barrels (dark-stippled). Some sAPF images for the putative 72 and 108mers have apparent six-fold radial symmetry. The small gray semi-ovals on the perimeter of the 6-fold schematics represent peripheral S1 segments. Circular Images shown for putative 60mers and 84mers may correspond to Type A models. The diameter, D_wall_, to the center of the wall was approximated by the diameter of B2 (0.19M) for tri β-barrel models and by the diameter half-way between B2 and B3 for tetra-β-barrel models (~0.19M/2 nm for 6-fold symmetric Type A models and 0.19(M + 6)/2 nm for 6-fold symmetric Type B models, where M is the number of monomers in the assembly). Note that the walls of the putative tri-β-barrels are considerably thinner than those of the putative tetra-β-barrels.

Our Type B models for sAPFs with 6-fold radial symmetry are more complex, but also more constrained. These models specify that all four β-barrels have S/N values of 1.0 and that each barrel has 12 more strands than the barrel it surrounds. This is made possible by the presence in each unit cell of an atypical monomer with an S1a-S1b β-hairpin in B4, a S2 β-strand in B3, and a S3 β-strand in B2, as illustrated in Fig. 9 and similar to the dodecamer model of Fig. 5b. The typical monomers of the outer barrels may have double U-turn conformations (S1b-S1a-S3 of Figs. 3g and 4c; S3 not shown in Fig. 3) with S1b strands forming an additional, but partial, antiparallel β-sheet layer and S1a segments fitting between S2-S3 linkers. If so, these S1b sheets will extend out from the outer barrel to form radially-symmetric nodules on the exterior of the sAPFs (Figs. 8 and 9). The structure of the entire assembly is determined by the unit cell (indicated by the dotted triangle in Fig. 8) and symmetry operations. For example, the structure with 6-fold symmetry can be generated by making a copy of the unit cell and rotating the copy by 180° about the Y axis. The assembly can be completed by rotating copies of the two antiparallel unit cells in increments of 60° about the Z axis.

For sAPFs with 6-fold radial symmetry and 2-fold vertical symmetry, Type A models are applicable only for APFs formed by an odd number of dodecamers and Type B models are applicable only for APFs formed by an even number of dodecamers: *i.e*., if there are 12 unit cells and Type A models have one more S3 β-strand in B3 than in B2, then the number of monomers per sAPF must be (1 + 2)12, (2 + 3)12, (3 + 4)12 ….; Type B models have the same number of S3 strands in B2 and B3 and thus the number of monomers per sAPF must be (1 + 1)12, (2 + 2)12, (3 + 3)12,…. These constraints may explain why 6-fold radial symmetry is not observable in some sAPFs (Type A structures) and is observable in other sAPFs (Type B structures).

If the number of S3 strands in B2 and B3 can be integer multiples of six in addition to a multiple of twelve (*e.g.*, 66 and 78), then sAPFs can belong to Type A regardless of the number of dodecamers in their precursors. When the number of monomers, M, is 132 or more, the S/N values of B2 and B3 can be 1.0 while S/N of B1 increases from 0.33 towards 1.0 and S/N for B4 decreases from 1.40 toward 1.0 as M increases, all while maintaining gap distances of ~ 1.0 nm between all barrels. Although this explanation may explain why few very large sAPFs exhibit radial symmetry, other possibilities cannot be excluded.

**Figure 9.**
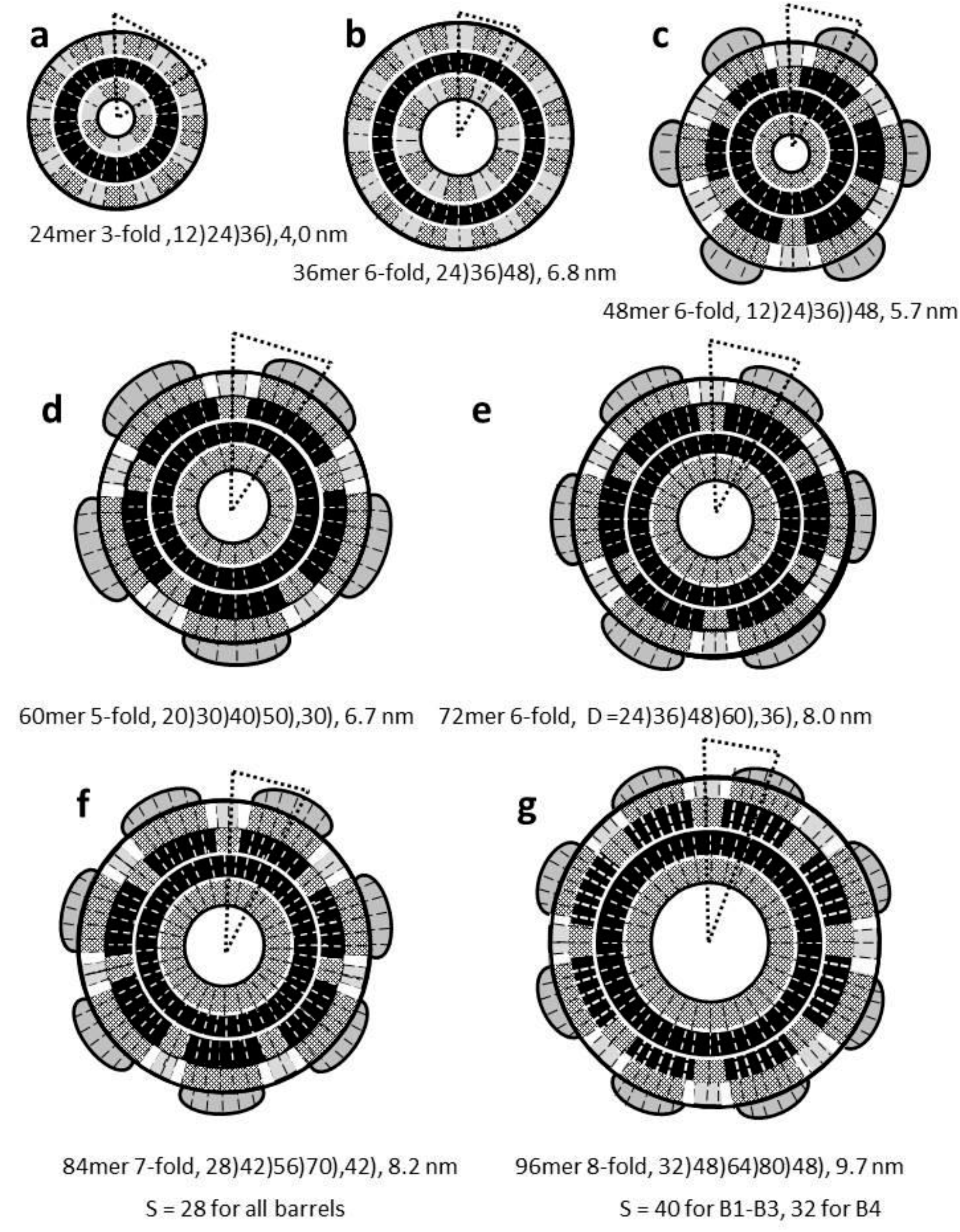
Schematic representations of tri-β-barrel and Type B tetra-β-barrel models of sAPFs. The shade coding is the same as in Fig. 5. All β-barrels have S/N values of 1.0 except for the last two models. Adjacent monomers are antiparallel, giving the assemblies 2-fold vertical symmetry. All assemblies have a multiple of 12 monomers; the number of strands in each barrel is listed below each schematic, followed by the diameter to the center of the wall (D_wall_, the center of B3 for tri-β-barrels and the border between B2 and B3 for tetra-β-barrels). All β-barrels have S/N values of 1.0 except for the last two models. Unit cells from which each assembly can be generated are indicated by dotted triangles. (a & b) Tri-β-barrel models composed of 24 and 36 Aβ42 monomers. (c-g) Type B models: Each unit cell contains an atypical monomer with a S1a-S1b β-hairpin in B4, a S3 β-strand in B3, and a S2 β-strand in B2; the unit cells are virtually identical for the last four models. Also, S1b (gray) and possibly S1a (not shown) strands may add one or two additional β-sheet layers peripheral to the S2 β-strands of B4. (c) A 48mer with 6-fold radial symmetry. (d) A 60mer model with 5-fold symmetry formed from five dodecamers. (e) A 72mer with 6-fold radial symmetry. (f and g) Models with 7-fold and 8-fold radial symmetry that are presumably formed from seven and eight dodecamers. S = 28 for all barrels of the 7-fold and S = 40 for B1-B3 and 32 for B4 of the 8fold models.

Fig. 10 shows numeours sAPF images with apparent 6-fold radial symmetry. D_wall_ values calculated for sAPFs with apparent 6-fold radial symmetry range from ~6 to ~17 nm (Fig. 10), which corresponds to Type B models with 48, 72, 96, 120, 144, and 168 monomers.

**Figure 10.**
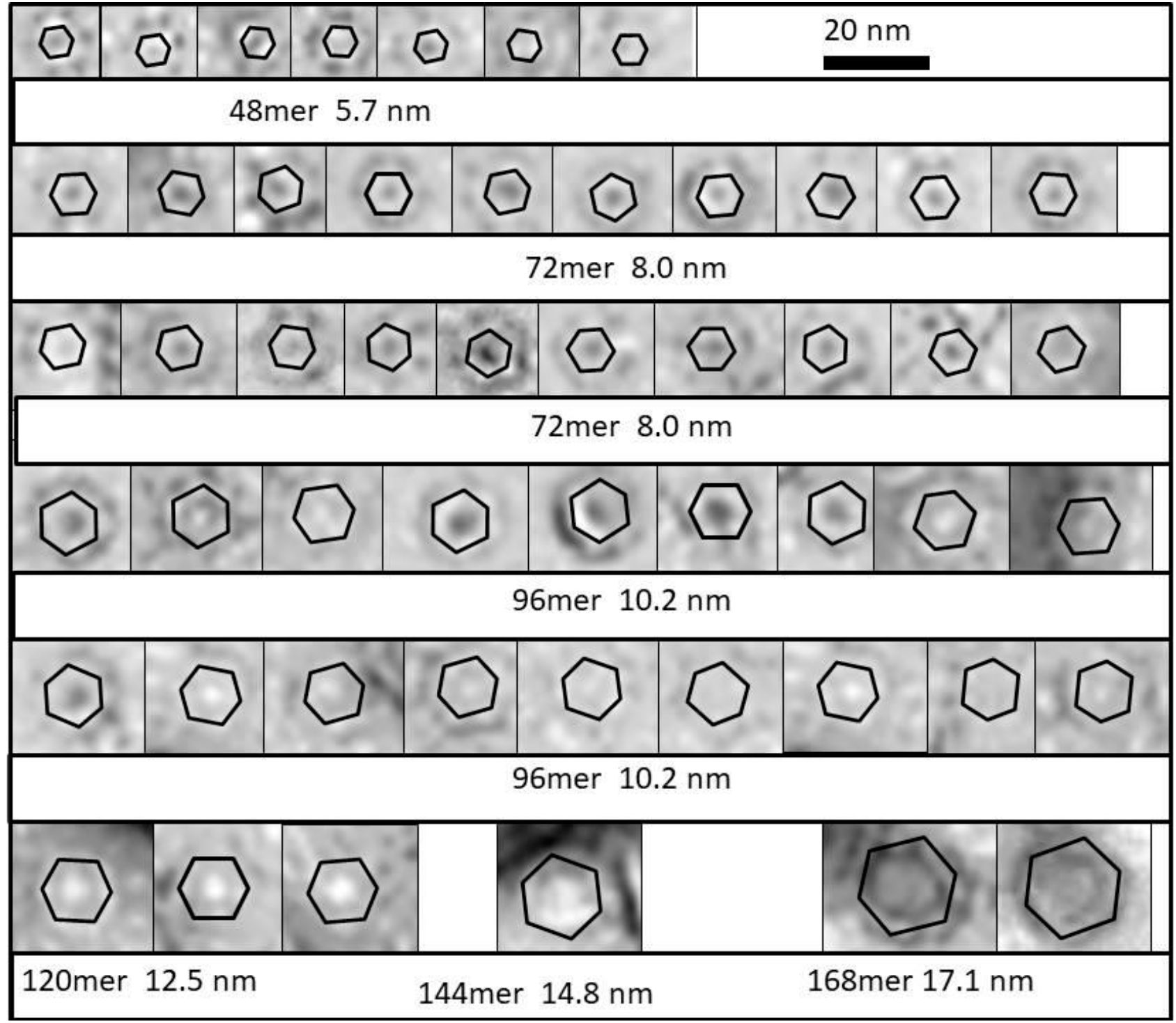
Images of sAPFs that exhibit, or exhibit features of, 6-fold radial symmetry. The predicted diameter to the center of the wall, D_wall_, illustrated by the hexagons, is listed below the images preceded by the number of monomers predicted by Type B models. Vertexes of these hexagons were positioned to correspond to dark spots on the perimeters of the sAPFs and/or sides of the hexagons that were positioned to align with apparently linear regions of the APF parimeters or sometimes with the inner edges of the walls.

Although sAPFs with apparent 6-fold radial symmetry are more common, some assemblies may retain the radial symmetry of their beaded precursors. A Type B sAPF may have 5-fold radial symmetry if it is formed from 5 or 10 dodecamers (Fig. 9d and Fig. 11). In these models, the number of strands in each successive barrel increases by ten, creating a gap distance between barrels of ~0.95 nm when S/N = 1.0, well within the limits. Likewise, a sAPF may have 7-fold and 8-fold radial symmetry if it is formed from 7 (or multiple of 7) or 8 dodecamers. In these cases the number of monomers in each successive barrel for Type B models increases by 14 or 16 (Fig. 9f and 9g and Fig. 11). If S/N = 1.0 the gap distance would be greater than we allow. However, radial symmetry can be achieved if S is the same for all four barrels of an assembly and/or a multiple of 14 or 16 for 7-fold and 8-fold models. The values of S listed in Fig. 11 were selected to produce gap distances between barrels of ~ 1.0 nm.

**Figure 11.**
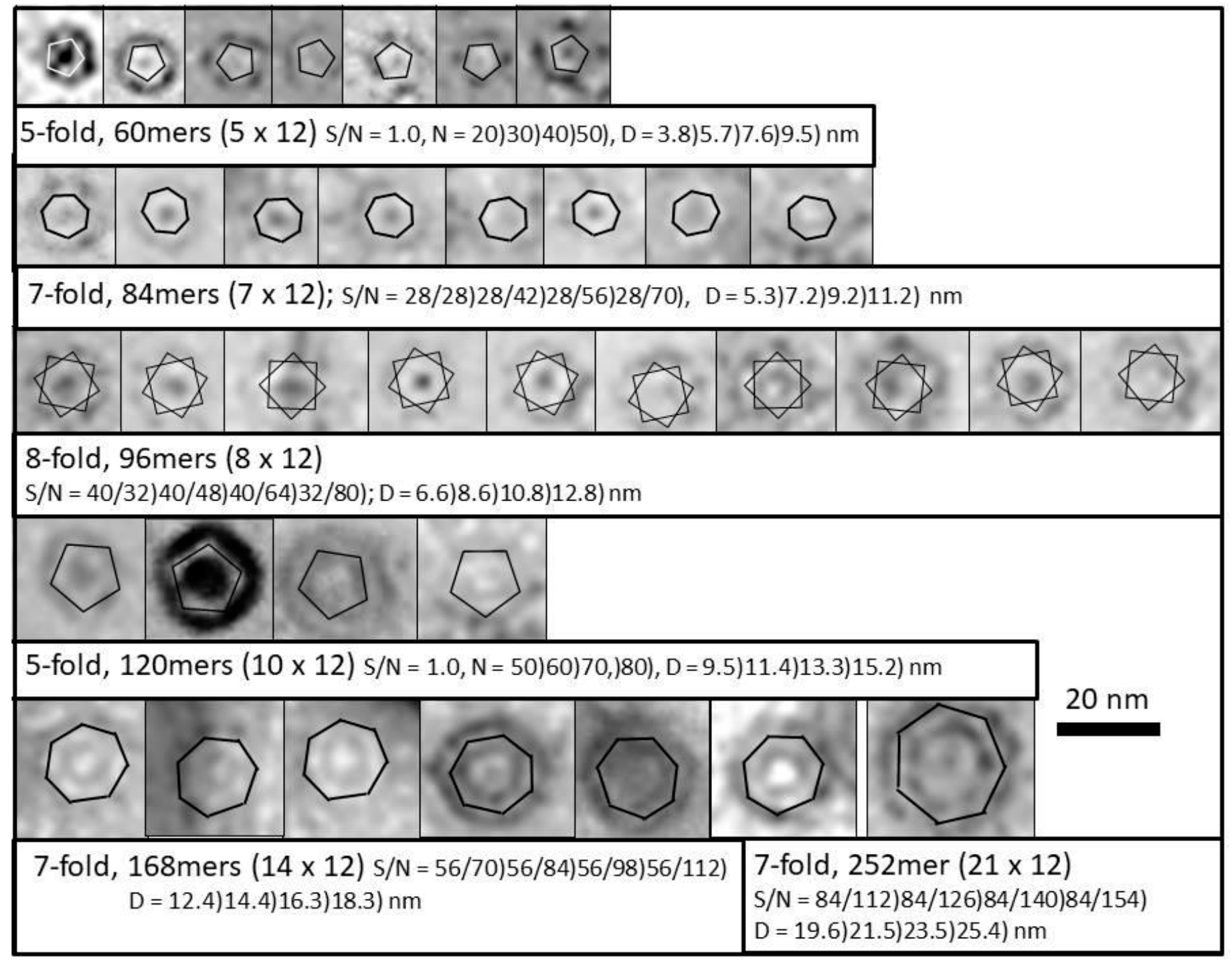
Images of sAPFs that exhibit, or may exhibit, 5-fold, 7-fold, and 8-fold radial symmetry. The diameters of the pentagons, heptagons, and 8-pointed stars approximate calculated values of D_wall_ (Fig. 9). The number of monomers, S/N values, and diameters of the four concentric barrels are listed below the images.

### Wall Thickness of Tetra-β-barrels

Some EM images of sAPFs with diameters greater than 12 nm show two concentric rings at the perimeter that we have measured to be about 4.9 nm apart regardless of the size of the APF (Fig. 12). We hypothesize that these rings correspond to the inner and outer edges of the β-barrel assemblies. If S/N = 1.0 for all barrels and each of four concentric β-barrels has 12 more strands than the barrel it surrounds, then the thickness of the wall should be (3)(1.14) + 1.0 = 4.4 nm (1.14 nm is the distance between each barrel and 1.0 nm is added for side chains), in good agreement with the measured distance. These results support our hypothesis that four concentric β-barrels form the walls of sAPFs. We cannot exclude the possibility, however, of a fifth barrel (add another 1.14 to reach 5.5 nm) since we are unsure how much side-chains contribute to the dimensions, and S1 segments could comprise an additional exterior β-barrel and interior or partial β-barrel.

**Figure 12.**
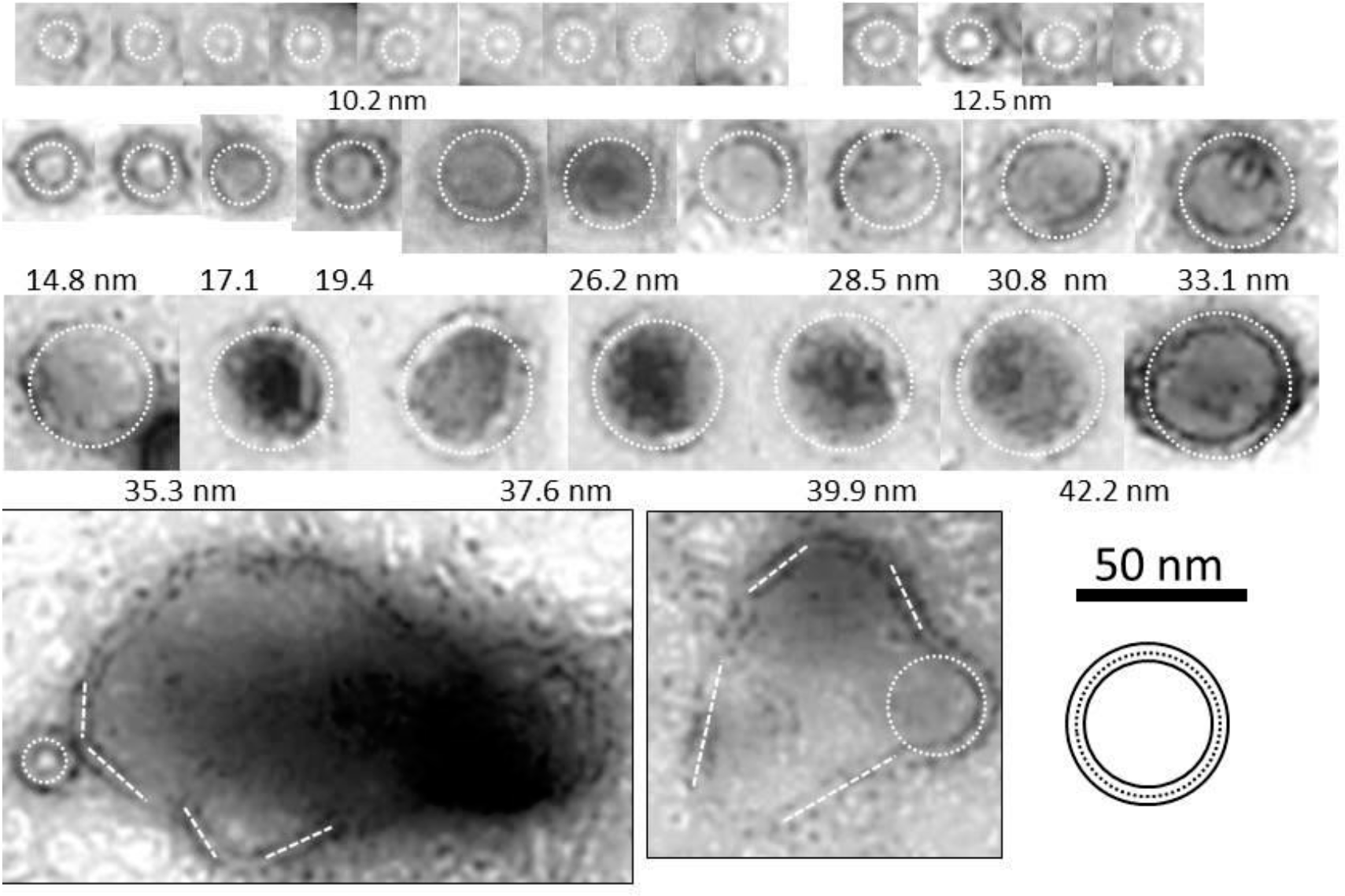
Images of sAPF with two concentric rings. The dashed circles positioned between the two rings indicate the dimensions to the center of the wall and have diameters predicted by our models beginning with a D_wall_ of 10.2 nm. The distance between the two rings was measured independent of the theory by using Microsoft PowerPoint 2010 to match each ring with a circle and then by calculating the radius of each circle. This averaged 4.9 nm, the distance between the solid circles outside and inside the dashed circles of the schematic on the bottom right. The two bottom images are for extremely large assemblies that have irregular shapes. Rectangles were superimposed on relatively straight portions of the walls to calculate the distance between the two rings around these gigantic sAPFs; which averaged the same as that between the rings of smaller sAPF.

## Part 4: Transmembrane Channels

Although APFs do not form transmembrane channels and are not highly toxic, they may have structures similar to β-barrel Aβ42 channels that form from Aβ42 oligomers ^10^. Here we expand and alter our previous models of Aβ42 channels formed from soluble Aβ42 hexamers ^18^.

### Microscopy images of membrane-imbedded Aβ42 assemblies

Our channel models have been influenced by freeze fracture EM images of transmembrane Aβ42 assemblies (original micrograph in Supplement Fig. S2). Fig. 13 shows enlargement of relatively circular isolated bodies segregated into eight categories and schematics of models that we propose are derived from two to nine hexamers.

**Figure 13.**
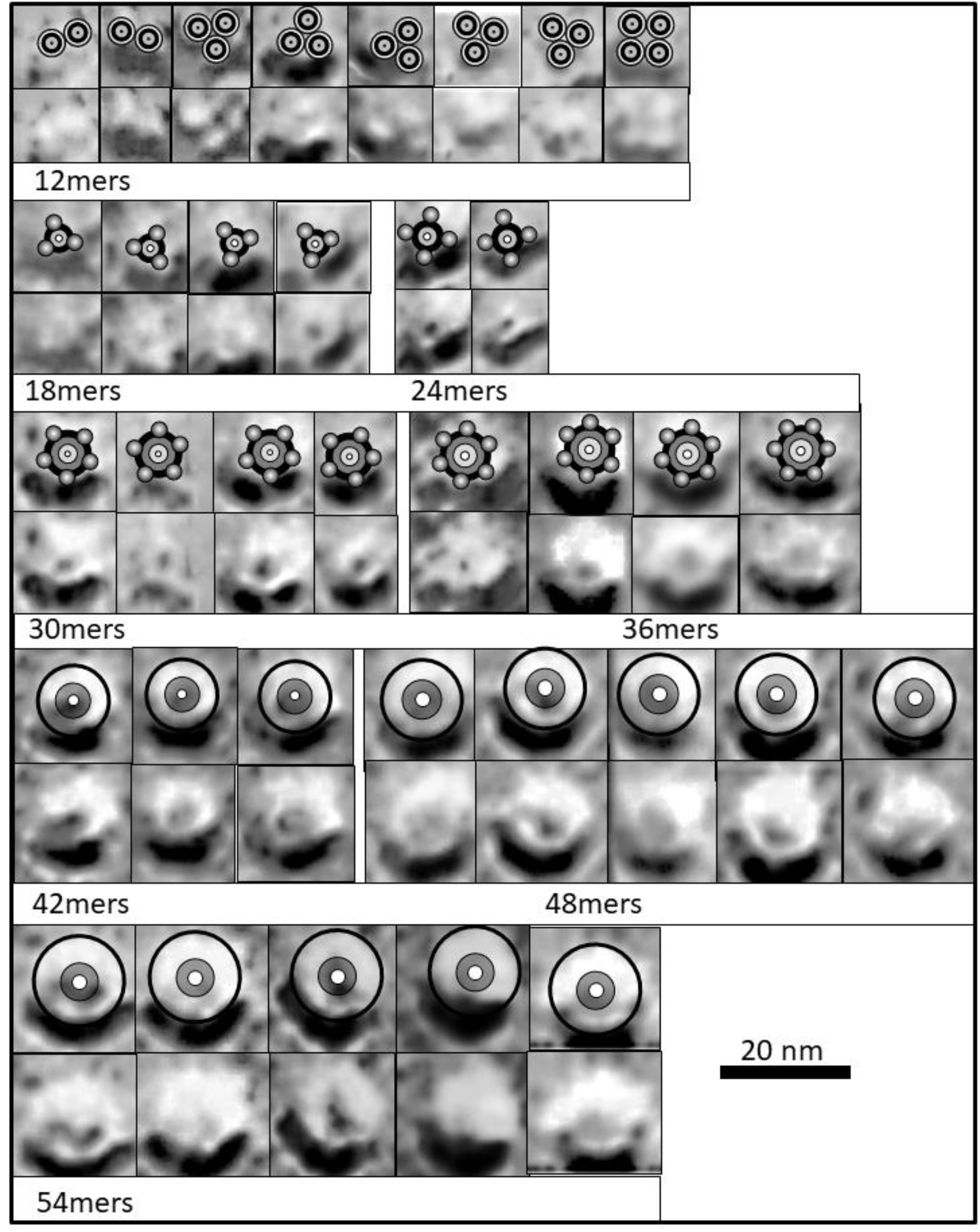
Freeze fracture image of membrane-embedded circular Aβ42 assemblies (from original by G. Zampighi and N. Arispi ^18^). The images are grouped according to how well their diameters and shapes correspond to the models presented below.

The freeze fracture studies of transmembrane structures have been complemented by atomic force microscopy analysis of membrane-embedded Aβ assemblies. These studies reveal clusters of three to six peaks that extends into the aqueous phase by about a nanometer ^37,38^. Unfortunately, the relatively large diameter of the probe’s tip (~ 60 nm) makes the lateral dimensions of these structures difficult to determine.

**Figure 14.**
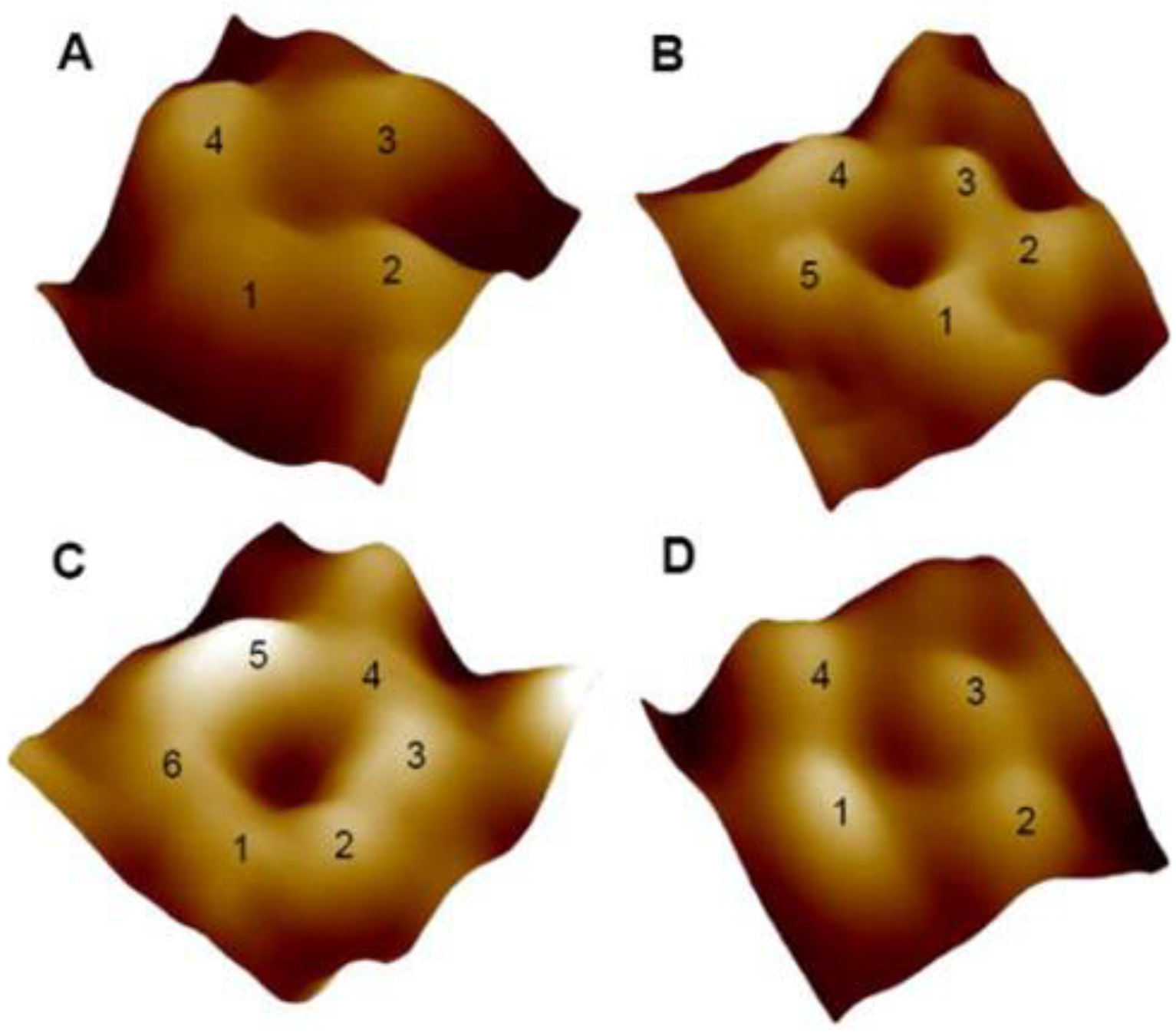
Atomic force microscopy images of Aβ42 assemblies. Image sizes are 11.5 nm for (A), 18.1 nm for (B), 13.2 nm for (C), and 14.4 nm for (D). Reproduced with permission from Connelly *et al.* Page 16 ^38^.

### Membrane Insertion Mechanisms

Before describing our models, we will consider how soluble Aβ oligomers could interact with and traverse a membrane to form channels. A multistage process is consistent with the time course of channel formation observed by Bode *et al.* ^12^: it takes a few minutes before channels start forming after neuronal membranes are exposed to Aβ42 oligomers and about ten minutes for half of the final number of channels to form. Fig. 15 illustrates three plausible multistage Aβ42 insertion mechanisms that have been simplified to convey general concepts rather than the likely highly dynamic and disordered insertion processes. The mechanism of Fig.15a produces vertically asymmetric channels (not proposed previously by us). If antiparallel β assemblies formed by S3 strands traverse the membrane, at least half of the S3 strands (those whose N-terminus end moves to the trans side of the membrane) must be accompanied by a S2 segment. Thus, a channel structure with N monomers could be formed in which an N-stranded antiparallel S3 β-barrel surrounds an N/2-stranded parallel S2 β-barrel. We call monomers with transmembrane S2 strands Pore-Lining (PL). S1 and S2 strands of the other monomers (called Aq monomers) remain in the aqueous phase on the cis side of the membrane. If the Stage 3 structure has N/2-fold radial symmetry, then there would be only two monomeric conformations (one for PL monomers and one for Aq monomers).

Alternatively, the β-barrels of the hexamers may split apart upon interaction with the membrane and associate to form raft-like assemblies that expose the hydrophobic S3 strands to the membrane’s alkyl phase. The structure depicted in Fig. 15b is similar to that of antiparallel Iowa-mutant fibrils ^6^ in which S2 β-sheets shield S3 β-sheets from water. S1a-S1b β-hairpins may occur at the ends of the rafts with their hydrophobic faces interacting with lipid alkyl chains. Hexameric rafts could interact side-by-side and end-on to form larger rafts with more extended β-sheets. These larger rafts could then fold into the membrane (like a clam closing) to form transmembrane β-barrels that are vertically symmetric or for which more than half of the S2 strands span the membrane. The transmembrane region could have an N-stranded S3 antiparallel β-barrel surrounding an N-stranded or N-1-stranded antiparallel S2 β-barrel.

The final possibility considered here is that the S3 β-barrels of hexamers remain intact when the hexamers traverse the membrane and none of the S1-S2 segments remain in the transmembrane region (see Fig. 15c). We call these Dumbbell structures because of their shape, *i.e*., the diameter of the aqueous S1-S2 domains on each end are likely to be greater than that of the six-stranded S3 barrel that connects them. If the insertion involves multiple hexamers or the hexamers aggregate after insertion, spaces between the S3 barrels would likely be filled with lipids and/or cholesterol. Thus, the transmembrane pore could be lined with lipid head-groups as has been proposed for some antimicrobial peptides ^39–41^.

**Figure 15.**
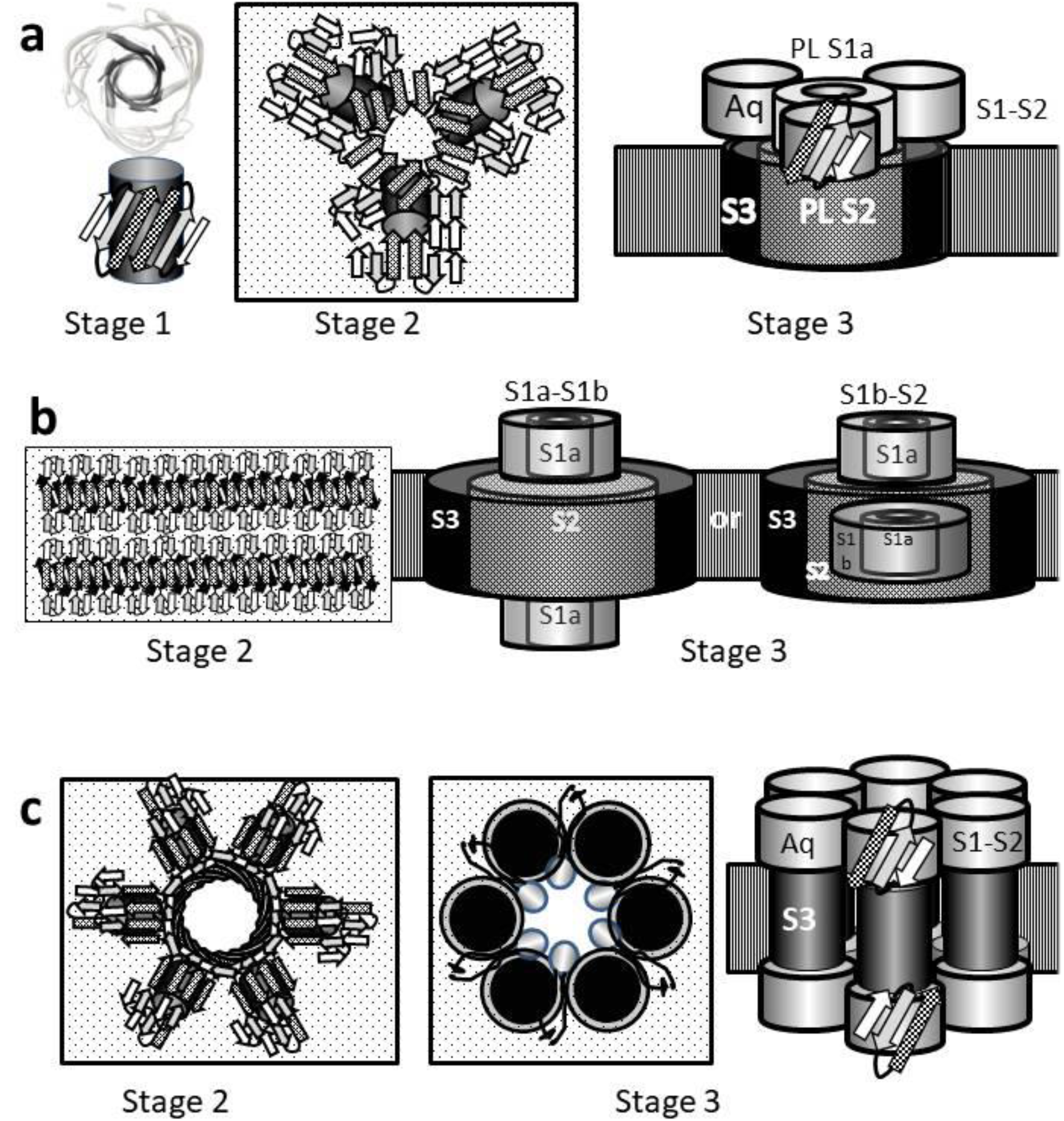
Multistage models of how soluble hexamers might interact with and then insert into a membrane to form channels. S1a strands are white, S1b strands are gray, S2 strands are stippled, and S3 strands or barrels are black. The lightly stippled rectangles represent the surface of the membrane and stripped blocks represent the hydrophobic portion of the membrane. (a) Formation of a vertically asymmetric 18mer channel. Radially outer barrels of the stage 3 model are transparent so that inner barrels can be seen. (b) Formation of a vertically symmetric channel in which all S2 strands span the membrane or asymmetric channel in which most S2 segments span the membrane. (c) An assembly of six hexamers with only 6-stranded S3 barrels in the transmembrane region and S1-S2 domains in the aqueous phases on both sides of the membrane. The egg-shaped images lining the pore represent lipid head groups and the black squiggly lines connected to them represent alkyl chains.

### Transmembrane concentric β-barrel theory and descriptions

We have placed the following constraints on our transmembrane models: (1) the monomers of a channel should have only a few different conformations (as reported by Serra-Batiste *et al.* ^10^), (2) the distance between the backbones of concentric β-barrels should be between 0.9 and 1.2 nm, (3) the diameters of the proposed structures should be consistent with the freeze fracture images and possibly the AFM images, (4) all major segments (S1a, S1b, S2, and S3) are β-strands that form portions of β-barrels, (5) all S3 strands are part of an antiparallel transmembrane β-barrel that has a S/N ratio between 0.0 and 2.0 and that surrounds a transmembrane S2 β-barrel, and (6) most hydrophobic side chains are in a hydrophobic environment while most hydrophilic side chains are exposed to water.

#### Aqueous domains

Hydrophilic and potentially flexible segments connecting S1a to S1b, S1b to S2, and S2 to S3 allow for multiple conformations (Fig. 3), especially for the aqueous domains. Nonetheless, S1a, S1b, and S3 likely comprise integral components of the assemblies and should not be simply ignored. An intriguing possibility for some aqueous domains is that these segments form a β-barrel (Fig. 16a) that is responsible for the peaks observed with AFM (Fig. 14). The interior of this putative β-barrel is filled with hydrophobic side-chains whereas the exterior is dominated by hydrophilic side-chains, most of which are charged (Fig 16b). An exterior region in the middle of S2 containing hydrophobic V18 and F20 side-chains is surrounded by charged side-chains and may correspond to points of contact between adjacent hexamers or dodecamers (Fig. 16c and Fig. 5e). If the barrel 9-stranded each hexamer has three such regions due to the 3-fold symmetry of the barrel. If these putative β-barrels form the aqueous domains of dumbbell structures, then contacts at these regions could result in formation of a hexagonal lattice (Fig. 16d).

**Figure 16.**
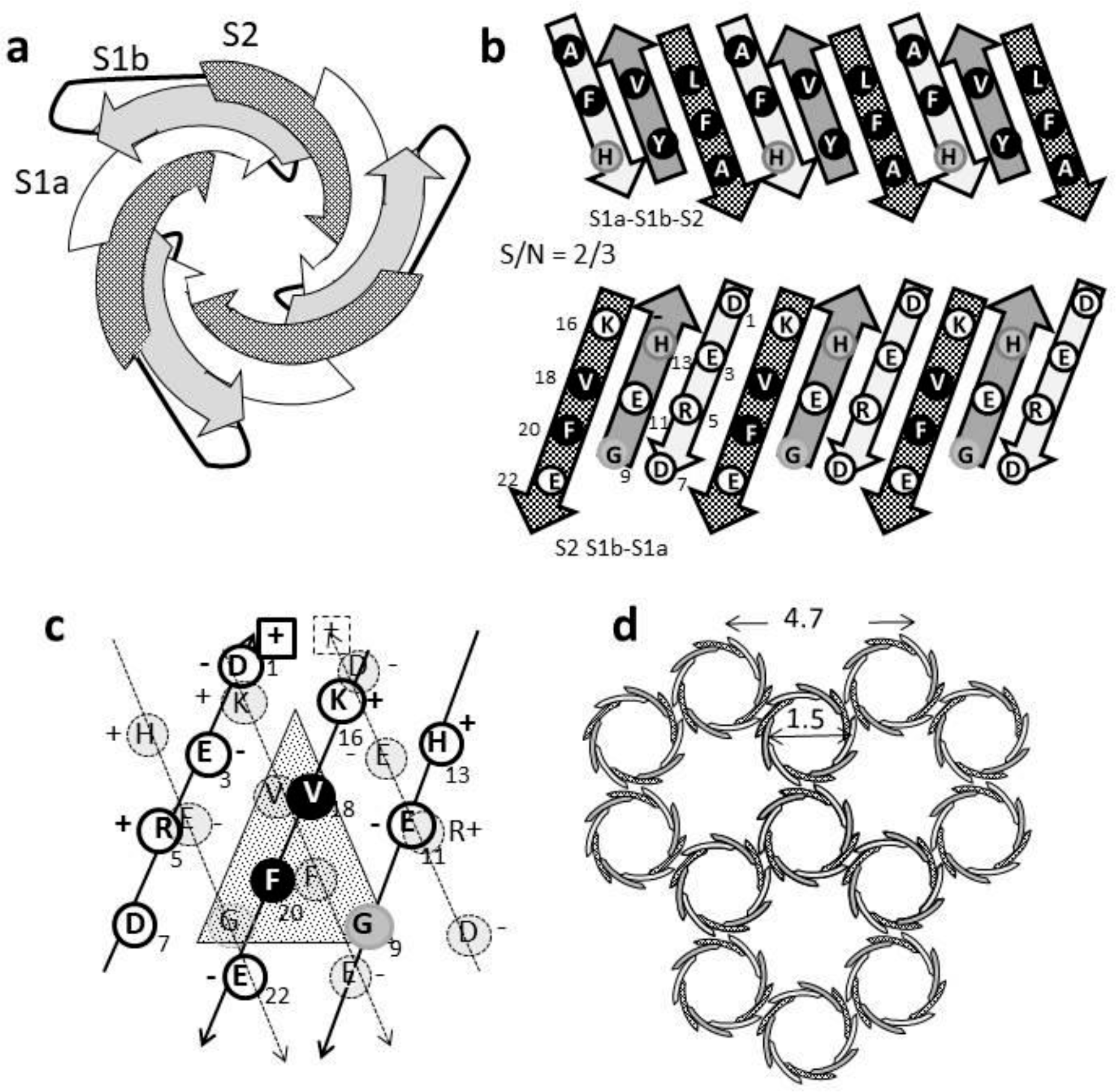
Nine-stranded β-barrel model of some aqueous domains formed by three S1a-S1b-S2 triple-stranded antiparallel β-sheets. (a) Schematic of the β-barrel as viewed from the top. (b) Repesentation of the side-chains on the interior (top) and exterior (bottom) of the barrel viewed as if the barrel were split open and spread flat. Hydrophobic side-chains are repesented by black circles and charged side-chains by white circles. (c) Representation of interactions between adjacent β-barrels in the region surrounding V18 and F20 in the middle of S2. The barrel on the far side is represented by heavier circles and solid arrows; the barrel on the near side by lighter dashed circles and lines. (d) A putative hexagonal lattice formed from the β-barrels with three radially symmetric interaction regions per barrel.

#### Conventional Channel Models

The models of Figs. 13 increase in increments of six monomers from a 12mer to a 54mer. The simplest and most constrained model is the 12mer (Fig. 17a). It is vertically asymmetric and has only two types of monomers (aqueous and pore-lining); giving each of the six unit cells of the transmembrane region only two S3 strands and one S2 strand. This requires S/N values of 2.0 for the antiparallel 12-stranded S3 β-barrel and the parallel six-stranded S2 β-barrel. The unit cell of the aqueous domain contains a triple-stranded Aq S1a-S1b-S2 β-sheet and a PL S1a strand from the PL monomer. This allows an 18-stranded S1a-S1b-S2 barrel (similar to the barrel described above but twice as large and with 6-fold radial symmetry) to surround a six-stranded S1a barrel with the hydrophobic side-chains of S1a oriented outwardly and interacting with the hydrophobic side-chains of the Aq barrel’s interior. The pore size through the pore-lining barrels of ~ 0.4 nm is so small that it might not conduct ions and or be visible in the freeze-fracture images. The diameter of the Aq domain is greater than that of the transmembrane region, and each Aq domain should contain six radially symmetric regions similar to those of Fig. 16d that could facilitate interactions among the 12mers. This may explain why so many of the putative 12mers of Fig. 13 cluster. The presence of exterior binding regions could also allow dumbbell assemblies to bind to 12mers, which could explain why some of the putative 12mer clusters appear to have additional material attached to the 12mers.

Monomers of unit cells of the 18mer and 24mer models have a similar transmembrane topology; however, the AFM and freeze fracture images are more consistent with the Aq S1a-S1b-S2 segments of each unit cell forming a distinct 8- or 9-stranded β-barrel. (The models of Fig. 17b have 8-stranded barrels with one Aq monomer contributing a S1a strand to the central S1a barrel, but the data are not sufficently precise to distinguish between the two possibilities.) Thus, the aqueous domain 18mers and 24mers would have three and four Aq β-barrels surrounding a 12- and 16-stranded β-barrel. The three Aq monomers of the unit cell would not have identical conformations; *i.e*., the differing symmetries for the Aq and transdomains require the S2-S3 linkers to have different conformations. The Aq β-barrels may be responsible for the presence of three or four apparent nodules on the perimeters of the putative 18mer and 24mer images of Fig. 13.

The increase in the sizes of the transmembrane S2 β-barrel for 30mers and 36mers should allow the PF S1a strands to reverse direction and form two β-turns and a β-hairpin in the aqueous domain while a formally Aq S1a-S1b-S2 segment enters the transmembrane region to form the narrowest part of the pore. The sizes and shapes of these models are consistent with the freezefracture images paired with these models in Fig. 13.

**Figure 17.**
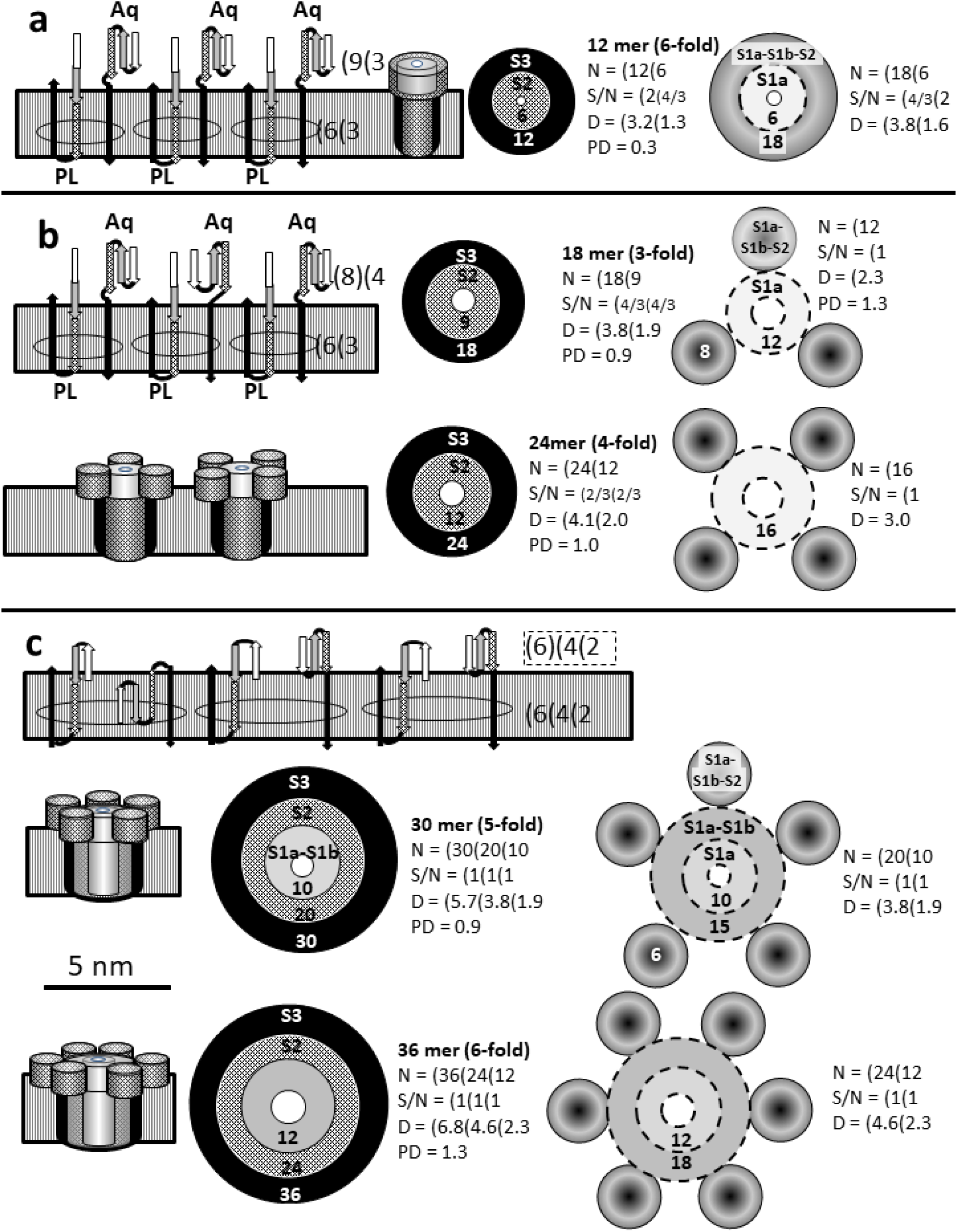
Schematics of small to medium size channels formed from two to six Aβ42 hexamers. Topologies of the hexameric unit cell strands are depicted on the upper left of each section (S3 strands are black arrows, S2 strands are stippled arrows, S1b strands are gray, and S1a strands are white). Closely spaced adjacent strands represent β-hairpins; those farther apart represent β-U-turns. The striped rectangles represent the transmembrane region. The number of strands in each barrel of the unit cell is listed on the right side with the ( indicating part of a β-barrel. A double () symbol indicates that the unit cell has an entire β-barrel. Aq and PL indicate the two types of monomers. Side views of the models are represented by cylinders with only the edges of the outer S3 barrel shown in black. Transmembrane concentric β-barrels of channel structure viewed through the pore are represented by black circles for S3, stippled circles for S2, and gray circles for S1a and S1b. The innermost white circle represents the pore. Aqueous domains are illustrated on the right (those with curved or donut surfaces represent S1a-S1b-S2 β-barrels, flat dark and light gray circles represent S1b and S1a β-barrels. The number of strands in the β-barrel is indicated at the bottom of each circle. Parameters of the assemblies are listed to the right of each schematic: *i.e*., the number of monomers in the assembly and its radial symmetry are listed first in bold; followed by N, the number of strands in each barrel; followed by the S/N ratio for each barrel; followed by the diameter, D, of the backbone of each barrel. PD strands for the estimated Pore Diameter, which is 1.0 nm less than the diameter of the smallest β-barrel of the aqueous domain. (a) A 12mer formed from two hexamers. This is the simplest structure with only two types of subunits repeated six times. The Aq S1a-S1b-S2 strands form an 18-stranded β-barrel that surrounds a 9-stranded β-barrel formed by S1a segments of the PL monomers. (b) Models of the 18mer and 24mer. The transmembrane topology is similar to that of the 12mer; however, the Aq S1a-S1b-S2 segments form an 8-stranded β-barrel within each hexameric repeat. These surround a β-barrel; each unit cell of this barrel has three S1a strands from the PL monomers and one S1a strand from an Aq monomer. (c) Models of 30mer and 36mer channels. The topologies of these structures differ because the S1a segments of the PL monomers have reversed directions to form S1a-S1b β-hairpins and U-turns and one of the former Aq S1a-S1b-S2 segments has entered the transmembrane region, and the Aq S1a-S1b-S2 barrels are 6-stranded with an S/N value of 4/3.

As more hexamers are added to the assemblies, the S3 β-barrel may become large enough to surround most of the S2 segments and/or allow some S1-S2 segments to move into the aqueous phase on the trans side of the membrane. The transmembrane topology of the models of Fig. 18a-d is similar to those of the smaller channels except for two monomers. The unit cell of TM regions of these models has six S3, five S2, and four S1a-S1b U-turns (Fig. 18a, see Fig. 3h for schematic of S1a-S1b-S2 double U-turn structure). The aqueous domain has one S2, two S1b, and two S1a strands; the unit cell of S1b and S2 strands comprises a β-barrel that surrounds a relatively small S1a β-barrel. Thus, the 42mer has a 21-stranded β-barrel around a 14-stranded β-barrel (the same as for Aerolysin ^26^) and the 54mer has a 27-stranded β-barrel around an 18 stranded β-barrel (the same as for Lysenin ^27^).

The outer diameters of the three largest categories of assemblies in Fig. 13 correspond closely to those of our models of 42mer, 48mer, and 54mer channels. A shadow appears on the lower left side of the tops of some of these assemblies. We propose that these shadows arise from the concentric S1b-S2 and S1a β-barrels that extend into the aqueous phases (represented by gray circles in Fig. 13).

**Figure 18.**
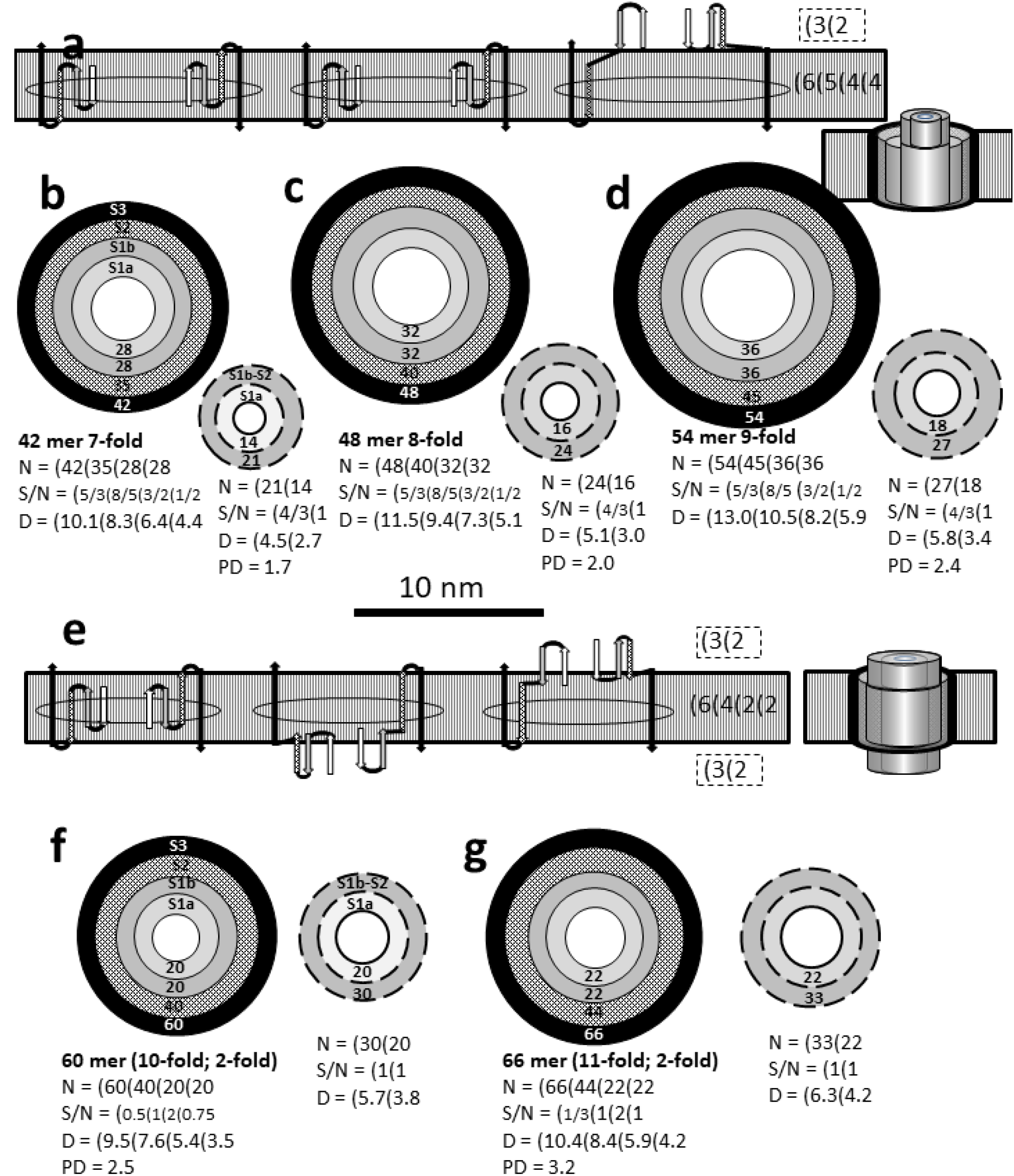
Schematic representations of 42mer, 48mer, 54mer, 60mer, and 66mer channel models. Structure parameters and images have the same meaning as in Fig. 17. (a) Transmembrane topology of a unit cell for vertically asymmetric models. (b-d) The four concentric circles on the left represent concentric transmembrane S3, S2, S1b, and S1a β-barrels. The circles with dashed lines on the lower right represent aqueous domains on the cis side of the membrane; β-barrels formed by S1b and S2 strands surround an S1a antiparallel β-barrel. (e) Transmembrane topology of a unit cell with 3 monomer conformations and 2-fold vertical symmetry. (f and g) Schematics for models with 60 and 66 monomers.

Some S1 and S2 segments of large assemblies may flip to the trans side of the membrane to create channels with 2-fold vertical symmetry, as illustrated in Fig. 18 e-g. The transmembrane unit cell of these models has six S3 strands, four S2 strands, two S1b strands and two S1a strands; *i.e*., one S2 strand and two S1a and S1b strands have moved to the trans side of the membrane relative to the previous models. Thus, these models have aqueous domains on both side of the membrane, each with the same topology as proposed for 42mer, 48mer, and 54mer models. This transfer reduces the diameter of the transmembrane region. The respective outer diameters of 60mer (10.5 nm) and 66mer (11.4 nm) models are slightly less and slightly greater than that of the 42mer (11.2 nm). Thus, it is difficult to distinguish among these models in the freeze-fracture images. However, the pore diameters, PDs, of these five models are virtually the same as sizes calculated by Bode *et al.* ^12^ from single channel conductances.

### Dumbbell hexagonal lattice assemblies

The models described cannot account for freeze-fracture images that have elongated or irregular shapes that contain two or three dark spots that are 4-5 nm apart (Fig. 19). This distance is too short for the spots to correspond to pores of any of the models described above. Earlier we suggested that interactions among the 9-stranded Aq S1a-S1b-S2 β-barrels of dumbbell-shaped hexamers at three hydrophobic patches on S2 strands could lead to toroidal channels composed of six hexamers, and that such assembles might expand to form a hexagonal lattice in which adjacent channels share two hexamers. A single channel of this type would have about the same diameter as proposed for the 24mer, and thus would be difficult to identify in the freeze fracture images, especially if additional dumbbell hexamers bind on the perimeter. However, a tell-tell pattern could be revealed if the assemblies have two or more pores that are about 4.4 nm apart, the distance between adjacent toroidal pores predicted by the hexagonal lattice model. Fig. 19 illustrates that such images exist and are fit well by the hexagonal lattice dumbbell models. A few assemblies appear to have elongated pores rather than distinct spots (bottom row of Fig. 19). These images can be fit by assuming that the dumbbell hexamers need to be in contact with lipid, and when the assemblies become sufficiently large, dumbbell hexamers that would be buried in the assemblies are simply missing, thus allowing the adjacent pores to merge into elongated clefts.

Almost all of these images have additional material not explained by a hexagonal lattice of dumbbell structures. Some of this material may be formed by 12mers. The outer surface of the aqueous domains of our 12mer models is composed of a S1a-S1b-S2 β-barrel similar to the soluble domains of the dumbbells except its circumference is twice as large and has six radially symmetric potential hydrophobic S2 binding sites. This suggests that up to six dumbbell hexamers could bind to the perimeter of a 12mer, possibly forming part of a lattice similar to that depicted at the bottom of Fig. 19. Also, as mentioned earlies, the 12 likely self-associate, as illustrated by the third lattice at the bottom of Fig. 19. The first two images of the last row in Fig. 19 support this hypothesis: *i.e*., they have three or four spherical bodies with the same appearance as the putative 12mers of Fig. 13 but that are surrounded by additional material to form a large assembly. The size and shapes of these assemblies can be fit well by a model for which surrounding dumbbells account for the additional material surrounding and separating the 12mers.

**Figure 19.**
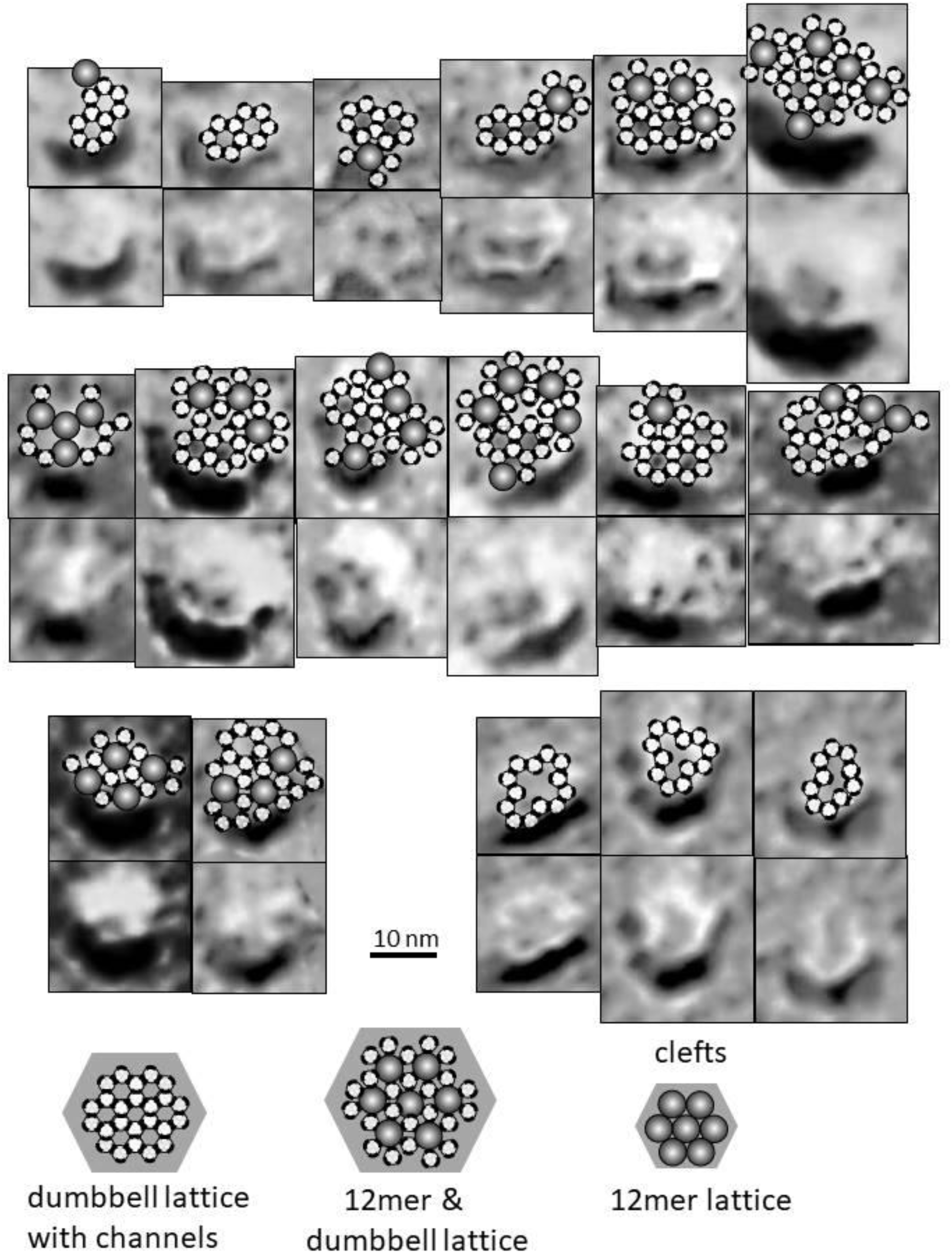
Non-circular images with multiple dark spots in freeze-fracture data (bottom rows) and dumbbell hexagonal lattice model predictions superimposed on the images (top rows). The small circles with diameters of 2.5 nm represent putative 9-stranded β-barrels formed by three sets of S1a-S1b-S2 strands. The black bars on the perimeters of the circles represent binding regions between hexamers. Regions in the center of the hexamer of hexamer rings represent toroidal pores and are positioned above the dark spots. The six-membered rings may not be complete in the interiors of assemblies we call clefts in the bottom row. Dumbbell structures may interact with 12mers represented by spheres. Plausible types of lattices formed by only dumbbells, by dumbbells and 12mers, and by only 12mers are illustrated at the bottom.

### Comparison of Pore Size Predicted by Single Channel Conduction to Models

Bode *et al.* ^12^ estimated Aβ42 pore diameters of 1.7, 2.1, and 2.4 nm, and possibly 2.6 and 3.2 nm based on an equation with the assumption that the pores are cylindrical with a uniform diameter throughout, and that ions diffuse through these nonselective pores in a manner similar to that of diffusion in water ^42,43^. Assuming that the pore diameter is about 1.0 nm less than the diameter of the wall for the narrowest pore-forming segments, the models of Fig. 18 predict pore diameters of 1.7, 2.0,2.4, 2.5, and 3.2 nm; virtually identical to the conductance-based approximations. Serra-Batiste *et al.* ^10^ estimated a smaller pore size of 0.7 nm; most models of Fig. 17 predict only slightly larger diameters of 0.9 (18mer and 30mer), 1.0 (24mer), and 1.3 nm (36mer). The dumbbell pores are likely to be less stable, and may be responsible for frequently observed flickering conductances.

## Part 5: EXPERIMENTAL TESTS

The models presented above should be considered as hypotheses to be tested experimentally. Experiments designed to answer the following questions are feasible: Do β-barrels comprise Aβ42 oligomers, APFs, and channels; if so, do these assemblies have concentric barrels; if so, how many barrels are there, are they radially and/or vertically symmetric, are they parallel or antiparallel, which segments form the barrels, how many strands do the barrels have, how much are the strands tilted, do β-barrel assemblies play an important role in AD, and can this knowledge lead to improved ways to prevent and/or treat AD?

### Single-Particle Cryo-Electron Microscopy

The ultimate test would be to determine the atomic structure of Aβ42 assemblies. The 2D EM data analyzed here were obtained over a decade ago and are limited by both quantity and quality. The fact that sAPFs and transmembrane Aβ assemblies can be classified by their sizes and shapes into specific categories suggests that modern methods such as single-particle cryo-electron microscopy (cryoEM) ^44^ that do not require crystal structures and that work exceptionally well for highly-symmetrical protein assemblies could be used to obtain much higher resolution data. For example, cryoEM has been used to determine near-atomic resolution structures of Aβ fibrils ^3^, toxins that contain concentric β-barrels ^26,27^, 3D images of two kinds of toxic amyloid α-synuclein oligomers that have pores through their centers ^45^ and are both present simultaneously, TRP channels ^46^, and potassium channels ^47^. It may be possible to use cryoEM to circumvent some polymorphic and crystallization problems that have hampered the field of Aβ assembly structure.

### NMR, EM, and Spectroscopy

Serra-Batiste, M. *et al.* ^10^ have apparently isolated a unique Aβ42 oligomer that, based on solid-state NMR analyses, contains β-barrels. With luck, they will be able to solve the 3-D structure of these oligomers, or at lease obtain sufficient data to determine which, if any, of the structures proposed here are viable candidates. It could also be useful to use either conventional or cryoEM methods to examine these assemblies in membranes, and Fourier transform infrared spectroscopy ^13^ to determine whether the β-barrels are parallel or antiparallel.

### Analogs that stabilize some structures while preventing others

Polymorphism and gradual transition from small to larger assemblies complicates structural studies of Aβ assemblies. It may be possible to avoid this problem by designing analogs that will stabilize one structure at the expense of others. For example, Lendel *et al.* ^48^ introduced two cysteines (one in S2 and one in S3) that caused S2 and S3 to form a β-hairpin. This linkage prevented formation of Aβ fibrils but not some protofibrils; allowing the 3-D structures of these assemblies to be analyzed with NMR. (Their data supported formation of hexamers that are dissimilar to the structures we propose).

While this particular linkage is incompatible with our models, others should be. In our models of hexamers, S1b and S2 form β-hairpins that do not form in fibrils (see Fig. 3) or in the transmembrane region of any of our channel models, other than in dumbbell assemblies. Thus, our models of hexamers, dodecamers and tri-β-barrel APFs might be stabilized by mutations of V12C + L17C or Y10C + F19C, whereas formation of fibrils, tetra-β-barrel sAPFs, and channels should be prevented. In some models S1a and S1b form a β-hairpin with F4 adjacent to V12 and H6 adjacent to Y10. Mutating these putative adjacent pairs to Cys should stabilize hexamers, the dodecamer of Fig. 5b, and dumbbell structures while precluding most others except possibly fibrils that have unresolved S1 segments.

Other tests include: (1) If three S1a-S1b-S2 segments (possibly stabilized by strategic disulfide bridges) form a 9-stranded β-barrel and if these barrels assemble into a hexagonal lattice, it may be possible to solve crystal structures of peptides with these sequences. Attempts to use antibodies to Aβ assemblies to treat AD have typically failed, perhaps because they did not bind to the toxic type of assembly. If S1a-S1b-S2 peptides form β-barrels, and these types of barrels are exposed in channels, would antibodies to these barrels be efficacious in treating AD? Could these structures be used as a vaccine against toxic forms of Aβ? (2) Interactions between Aβ42 hexamers that lead to formation of dodecamers and dumbbell lattices and interactions between dodecamers that lead to APFs may involve interactions between the exposed faces of antiparallel pairs of S2 segments; specifically K16, V18, F20, and E22. If so, mutations of V18 and F20 to hydrophilic residues (*e.g*., V18E and F20K) should allow formation of the hexamers but might inhibit formation of the larger oligomers. These mutations should also inhibit structures that contain S1b-S2 U-turns, such as Type B sAPFs and the 42-54mer channels. (3) Likewise, mutating hydrophobic residues of S1a (A2 and F4) and/or those of S1b (Y10 andV12) to hydrophilic residues should inhibit formation of assemblies in which these residues are buried *e.g*., hexamers, S1a-S1b-S2 β-barrels, and all channels. (4) The C-termini of three monomers are in close proximity in our hexamer models. Tethering these termini to a ligand with 3-fold symmetry might stabilize the antiparallel hexamer while preventing formation of other assemblies.

### Mutations

Numerous natural and created mutations affect Aβ toxicity and AD (see ALZFORUM site (https://www.alzforum.org/mutations/app) for a partial list). The Iowa mutation (D23N) responsible for some forms of early onset AD increases formation of antiparallel fibrils; perhaps because D23 forms a salt bridge with K28 of an adjacent subunit in parallel fibrils but not in antiparallel fibrils ^6^. Effects of the Osaka mutation (E22Δ) may be due to a similar effect since it would move the negatively charged D23 side chain from the hydrophobic side of S2 to the polar side, and move the hydrophobic V24 side chain from the polar side to the more hydrophobic side. Aβ42 mutations that increase the toxicity in a yeast screening system also increase formation of antiparallel assemblies ^14^. However, effects of most Aβ mutations that affect AD on formation and properties of Aβ oligomers, APFs and ion channels have not been studied.

Several familial mutations occur at E22 (E22K (Italian), E22G (Arctic), E22Q (Dutch)). E22 is positioned on the aqueous-exposed surfaces of fibrils and interacts with K16 of adjacent monomers in our antiparallel models. High pressure NMR of Aβ42 assemblies with E22G and D23N mutations revealed conformational perturbations with high pressure sensitivity at Q15, K16, and L17 ^49^.

### S1 Segment, Channel Selectivity and Channel Inhibitors and Blockers

Although the S1 segment is often ignored, it is an integral component of most of our models. Numerous mutations in the S1 segment affect toxicity and AD. A2T (Icelandic) is protective, whereas A2V is pathogenic. D7 at the end of S1a appears to be a hot spot: D7R (Taiwanese) and D7N (Tottori) are pathogenic. Three mutations (R5G, Y10F and H13R) in the rodent S1 segments dramatically reduce the toxicity of Aβ ^50^. Will these mutations inhibit formation of hexamers and/or channels? Zn^2+^ binds to and alters the structure of the N-terminus S1 region ^51^ and inhibits Aβ channels in lipid bilayers ^52^. Will Zn^2+^ also inhibit formation of Aβ42 channels in excised patches?

N-terminus truncation variants 1-x, 3-x, 11-x, and 17-x have been assessed in human amyloid plaques, and N-terminal truncation (probably 17x) appears to be involved in early amyloid pathology in Down’s Syndrome ^53^. Synthesized 17x Aβ peptides form channels in lipid bilayers ^54^, but it is not known whether they form channels in excised neuronal patches, and if so whether these channels have the same conductance properties as WT Aβ42 channels.

The polar face of S1a consists entirely of charged side-chains (D1, E3, R5, and D7) and forms at least part of the lining of the pore in all the conventional channel models presented here. If so, a R5E mutation should increase the permeability of cations and decrease it for anions. Likewise, D1K, E3K, and D7K mutations should increase permeation of anions and decrease it for cations.

Several molecules or heavy ions have been reported to block Aβ-channels in bilayers ^55,56^. Will any of these also block Aβ42 channels in excised patches? If so, will any of the mutations described above affect binding of these inhibitors?

### Receptor Binding and All-d-enantiomers

Numerous groups contend that Aβ oligomers affect neurons by binding to specific receptors. Bode *et al.* ^12^ reported that Aβ42 oligomers did not form channels in a substantial number of excised neuronal patches, suggesting that something other than a pure lipid bilayer is required: *e.g*., lipid rafts or a receptor. If a protein receptor is required for channel formation, then Aβ42 oligomers composed of an all-d-enantiomer and that forms channels in lipid bilayers ^57^ should not form the same types of channels as normal Aβ42 oligomers in excised patches.

### Atomic Scale Modeling and Molecular Dynamics Simulations

Molecular dynamic simulations in the absence of a preconceived model can sometimes be informative for relatively small peptides and small assemblies. For example, Sun *et al.* ^24^ found that in some simulation runs beginning with randomized peptides with the sequence of Aβ 16-22 (the core of S2) a six-stranded antiparallel β-barrel formed, and Qian *et al.* ^25^ obtained similar results for peptides with a Aβ_30-36_ sequence of S3. But these types of simulations last only a fraction of a second, and it is unrealistic to expect them to produce accurate predictions for larger peptides and larger assemblies that form gradually and often depend upon an initial seed structure. MD simulations of large assemblies performed on highly flawed initial models are likely to be meaningless, especially if the initial models are inconsistent with the basic β-barrel theory for concentric β-barrels composed of identical strands. Development of atomically explicit β-barrel models such as those we published previously ^17,18^ are simplified greatly by first predicting the number of monomers in the assemblies, the topology of the strands, the sheer number of the β-barrels, whether the strands are parallel or antiparallel, the symmetry of the assembly, and the positions of concentric β-barrels with respect to each other. Our new models presented here provide that information and thus can serve as a starting point for developing new atomically explicit models.

Nonetheless, we doubt that MD simulations are as informative as many believe. MD simulations of our earlier atomic scale models indicated that the antiparallel S3 barrels with 6 or 36 strands and S/N values of 1.0 remained extremely stable and maintain backbone hydrogen bonding between strands whether the assemblies are in water and shielded by S1 and/or S2 strands or span a lipid bilayer ^17,18^. However, MD simulations can be misleading and errors in some regions such as the S1 segments or the exclusion of some portions could introduce instabilities. Also, exclusion of important components, such a hexane in APF models and lipids or cholesterol in channels ^58^, could introduce apparent instability even if the model is otherwise correct. We have considered scores of alternative β-barrel models; only the simplest and most data-consistent of which were presented here. We doubt that MD simulations would help in distinguishing among these models. Thus, it seems prudent to wait until more precise structural data are available before attempting extensive atomic scale modeling of the large range of APFs and channel structures proposed here.

## Part 6: SUMMARY AND CONCLUSIONS

“Everything should be made as simple as possible, but not simpler.”

Albert Einstein, https://www.brainyquote.com/authors/albert_einstein

The microscopy data for membrane-bound structures and multiple channel conductances reported by Bode *et al.* ^12^ indicate that we should not think in terms of single structure. Although we have proposed more models than we would have preferred, we have proposed only as many as required to explain the available data.

Many may find the concept of tetra-β-barrels and/or gigantic β-barrels with diameters up to 75 nm difficult to accept, but the hypothesis is not as radical as it may appear. Both parallel and antiparallel Aβ fibril structures have been determined in which two S3 β-sheets are sandwiched between two S2 β-sheets ^1,2,6^. All of the interior monomers of these fibril structures have identical U-shaped conformations and identical interactions with neighboring monomers. The sAPF tetra-β-barrel structures we propose are similar: almost all monomers have the classic U-shaped S2-S3 β-structure, two hydrophobic S3 β-structures pack back-to-back, and these are sandwiched between two S2 β-structures. The major difference is that the β-barrel structures we propose for sAPFs are circular barrels instead of linear sheets as in fibrils.

Smooth APFs may have many features in common with channels such as concentric antiparallel β-barrels composed of S1, S2, and S3 segments that develop from soluble hexamers and similar arrangements of S1a, S1b, and S2 segments. The hypothesis that Aβ42 oligomers have structures similar to those of antiparallel β-barrel channel toxins is strengthened by findings that some antibodies recognize both types of structures ^15^ and by Fourier transform infrared spectroscopy studies indicating that Aβ42 oligomers have antiparallel β-structures remarkably similar to that of bacterial outer membrane porins ^13^. Thus, determination of APF structures could be informative about Aβ42 channel structures, concentric β-barrel structures, and how Aβ42 oligomers can form larger β-barrel structures.

But the ultimate purpose of this work is not to create molecular models or even to experimentally solve the structures of Aβ assemblies; it is to assist in improving prevention and treatment of AD. Development of improved prevention, treatments, and cures of this devastating and expensive disease may depend upon improved understanding of the underlying molecular structures and the processes that lead to their creation. We only hope that our analyses of the Aβ assembly process and the structures of these assemblies will help achieve that goal. The major point of our analysis is that the possibility of Aβ42 forming antiparallel β-barrels should be taken seriously and studied more extensively because they may contribute strongly to AD. If our basic concept that these assemblies have well-ordered, relatively simple β-barrel structures is valid, additional structural studies are likely to be productive.

## Authors’ contributions

Most of the theory and all figures were developed by HRG, who wrote most of the text. SRD performed the calculations for β-barrel parameters in Table S1 and provided valuable suggestions for the theory, figures, and editing of the manuscript. RK obtained and provided the EM data of the APFs and information regarding the images.

The authors have no competing interests.

Original EM images will be supplied by HRG at upon request.

## Acknowledgement

We thank Nelson Arispe for supplying the original freeze fracture image.

## References

1. Lührs T. et al. 3D structure of Alzheimer’s amyloid-β(1-42) fibrils. Proceedings of the National Academy of Sciences of the United States of America 102, 17342–17347, doi:10.1073/pnas.0506723102 (2005).

2. Petkova, A. T., Yau, W. M. & Tycko, R. Experimental constraints on quaternary structure in Alzheimer’s beta-amyloid fibrils. Biochemistry 45, 498–512, doi:10.1021/bi051952q (2006).

3. Gremer L. et al. Fibril structure of amyloid-β(1-42) by cryo-electron microscopy. Science 358, 116–119, doi:10.1126/science.aao2825 (2017).

4. Lu J. X. et al. Molecular structure of beta-amyloid fibrils in Alzheimer’s disease brain tissue. Cell 154, 1257–1268, doi:10.1016/j.cell.2013.08.035 (2013).

5. Soldner, C. A., Sticht, H. & Horn, A. H. C. Role of the N-terminus for the stability of an amyloid-beta fibril with three-fold symmetry. PLoS One 12, e0186347, doi:10.1371/journal.pone.0186347 (2017).

6. Qiang, W., Yau, W. M., Luo, Y., Mattson, M. P. & Tycko, R. Antiparallel beta-sheet architecture in Iowa-mutant beta-amyloid fibrils. Proc Natl Acad Sci U S A 109, 4443–4448, doi:10.1073/pnas.1111305109 (2012).

7. Mroczko, B., Groblewska, M., Litman-Zawadzka, A., Kornhuber, J. & Lewczuk, P. Amyloid β oligomers (AβOs) in Alzheimer’s disease. Journal of Neural Transmission 125, 177–191, doi:10.1007/s00702-017-1820-x (2018).

8. Fu L. et al. Comparison of neurotoxicity of different aggregated forms of Aβ40, Aβ42 and Aβ43 in cell cultures. Journal of Peptide Science 23, 245–251, doi:10.1002/psc.2975 (2017).

9. Arispe, N., Rojas, E. & Pollard, H. B. Alzheimer disease amyloid beta protein forms calcium channels in bilayer membranes: blockade by tromethamine and aluminum. Proc Natl Acad Sci U S A 90, 567–571 (1993).

10. Serra-Batiste M. et al. Aβ42 assembles into specific β-barrel pore-forming oligomers in membrane-mimicking environments. Proceedings of the National Academy of Sciences 113, 10866–10871, doi:10.1073/pnas.1605104113 (2016).

11. Serra-Batiste, M., Tolchard, J., Giusti, F., Zoonens, M. & Carulla, N. Stabilization of a Membrane-Associated Amyloid-β Oligomer for Its Validation in Alzheimer’s Disease. Frontiers in Molecular Biosciences 5, doi:10.3389/fmolb.2018.00038 (2018).

12. Bode, D. C., Baker, M. D. & Viles, J. H. Ion Channel Formation by Amyloid-β42 Oligomers but Not Amyloid-β40 in Cellular Membranes. Journal of Biological Chemistry 292, 1404–1413, doi:10.1074/jbc.M116.762526 (2017).

13. Cerf E. et al. Antiparallel beta-sheet: a signature structure of the oligomeric amyloid beta-peptide. Biochem J 421, 415–423, doi:10.1042/BJ20090379 (2009).

14. Vignaud H. et al. A Structure-Toxicity Study of Aß42 Reveals a New Anti-Parallel Aggregation Pathway. PLOS ONE 8, e80262, doi:10.1371/journal.pone.0080262 (2013).

15. Yoshiike, Y., Kayed, R., Milton, S. C., Takashima, A. & Glabe, C. G. Pore-forming proteins share structural and functional homology with amyloid oligomers. Neuromolecular medicine 9, 270-275 (2007).

16. Hu Z.-W. et al. Phosphorylation at Ser8 as an Intrinsic Regulatory Switch to Regulate the Morphologies and Structures of Alzheimer’s 40-residue β-Amyloid (Aβ40) Fibrils. Journal of Biological Chemistry 292, 2611–2623, doi:10.1074/jbc.M116.757179 (2017).

17. Shafrir, Y., Durell, S. R., Anishkin, A. & Guy, H. R. Beta-barrel models of soluble amyloid beta oligomers and annular protofibrils. Proteins 78, 3458–3472, doi:10.1002/prot.22832 (2010).

18. Shafrir, Y., Durell, S., Arispe, N. & Guy, H. R. Models of membrane-bound Alzheimer’s Abeta peptide assemblies. Proteins 78, 3473–3487, doi:10.1002/prot.22853 (2010).

19. Yun, S. J., Yun, S. J. & Guy, H. R. Analysis of the stabilities of hexameric amyloid-beta(1-42) models using discrete molecular dynamics simulations. J Mol Graph Model 29, 657–662, doi:10.1016/j.jmgm.2010.11.008 (2011).

20. Laganowsky A. et al. Atomic view of a toxic amyloid small oligomer. Science 335, 1228–1231, doi:10.1126/science.1213151 (2012).

21. Do T. D. et al. Amyloid β-Protein C-Terminal Fragments: Formation of Cylindrins and β-Barrels. Journal of the American Chemical Society 138, 549–557, doi:10.1021/jacs.5b09536 (2016).

22. Harmeier, A. et al. Role of amyloid-beta glycine 33 in oligomerization, toxicity, and neuronal plasticity. J Neurosci 29, 7582–7590, doi:10.1523/JNEUROSCI.1336-09.2009 (2009).

23. Bitan G. et al. A molecular switch in amyloid assembly: Met(35) and amyloid beta-protein oligomerization. Journal of the American Chemical Society 125, 15359–15365, doi:10.1021/ja0349296 (2003).

24. Sun, Y., Ge, X., Xing, Y., Wang, B. & Ding, F. beta-barrel Oligomers as Common Intermediates of Peptides Self-Assembling into Cross-beta Aggregates. Sci Rep 8, 10353, doi:10.1038/s41598-018-28649-7 (2018).

25. Qian, Z. Y., Zhang, Q. W., Liu, Y. & Chen, P. J. Assemblies of amyloid-beta(30-36) hexamer and its G33V/L34T mutants by replica-exchange molecular dynamics simulation. Plos One 12, doi:ARTN e0188794/journal.pone.0188794 (2017).

26. Iacovache I. et al. Cryo-EM structure of aerolysin variants reveals a novel protein fold and the pore-formation process. Nat Commun 7, 12062, doi:10.1038/ncomms12062 (2016).

27. Bokori-Brown M. et al. Cryo-EM structure of lysenin pore elucidates membrane insertion by an aerolysin family protein. Nat Commun 7, 11293, doi:10.1038/ncomms11293 (2016).

28. Murzin, A. G., Lesk, A. M. & Chothia, C. Principles determining the structure of beta-sheet barrels in proteins. I. A theoretical analysis. J Mol Biol 236, 1369–1381 (1994).

29. Chou, K. C., Carlacci, L. & Maggiora, G. G. Conformational and Geometrical Properties of Idealized Beta-Barrels in Proteins. Journal of Molecular Biology 213, 315–326, doi:Doi 10.1016/S0022-2836(05)80193-7 (1990).

30. Reboul, C. F., Mahmood, K., Whisstock, J. C. & Dunstone, M. A. Predicting giant transmembrane β-barrel architecture. Bioinformatics 28, 1299–1302, doi:10.1093/bioinformatics/bts152 (2012).

31. Zeytuni N. et al. Near-atomic resolution cryoelectron microscopy structure of the 30-fold homooligomeric SpoIIIAG channel essential to spore formation in <em>Bacillus subtilis</em>. Proceedings of the National Academy of Sciences 114, E7073–E7081, doi:10.1073/pnas.1704310114 (2017).

32. Schulz G. E. The dominance of symmetry in the evolution of homo-oligomeric proteins. J Mol Biol 395, 834–843, doi:10.1016/j.jmb.2009.10.044 (2010).

33. Goodsell D. S. & Olson A. J. Structural symmetry and protein function. Annu Rev Biophys Biomol Struct 29, 105–153, doi:10.1146/annurev.biophys.29.1.105 (2000).

34. Dean, D. N., Rana, P., Campbell, R. P., Ghosh, P. & Rangachari, V. Propagation of an Abeta Dodecamer Strain Involves a Three-Step Mechanism and a Key Intermediate. Biophys J 114, 539–549, doi:10.1016/j.bpj.2017.11.3778 (2018).

35. Kayed R. et al. Annular Protofibrils Are a Structurally and Functionally Distinct Type of Amyloid Oligomer. Journal of Biological Chemistry 284, 4230–4237, doi:10.1074/jbc.M808591200 (2009).

36. Huang D. et al. Antiparallel β-sheet structure within the C-terminal region of 42-residue Alzheimer’s β-amyloid peptides when they form 150 kDa oligomers. Journal of molecular biology 427, 2319–2328, doi:10.1016/j.jmb.2015.04.004 (2015).

37. Quist A. et al. Amyloid ion channels: a common structural link for protein-misfolding disease. Proc Natl Acad Sci U S A 102, 10427–10432, doi:10.1073/pnas.0502066102 (2005).

38. Connelly L. et al. Atomic force microscopy and MD simulations reveal pore-like structures of all-D-enantiomer of Alzheimer’s beta-amyloid peptide: relevance to the ion channel mechanism of AD pathology. J Phys Chem B 116, 1728–1735, doi:10.1021/jp2108126 (2012).

39. Cruciani R. A. et al. Magainin 2, a natural antibiotic from frog skin, forms ion channels in lipid bilayer membranes. Eur J Pharmacol 226, 287–296 (1992).

40. Durell, S. R., Raghunathan, G. & Guy, H. R. Modeling the ion channel structure of cecropin. Biophys J 63, 1623–1631, doi:10.1016/S0006-3495(92)81730-7 (1992).

41. Lipkin R. & Lazaridis T. Computational studies of peptide-induced membrane pore formation. Philos Trans R Soc Lond B Biol Sci 372, doi:10.1098/rstb.2016.0219 (2017).

42. Hille B. Pharmacological Modifications of the Sodium Channels of Frog Nerve. The Journal of General Physiology 51, 199–219 (1968).

43. Cruickshank, C. C., Minchin, R. F., Le Dain, A. C. & Martinac, B. Estimation of the pore size of the large-conductance mechanosensitive ion channel of Escherichia coli. Biophysical Journal 73, 1925–1931 (1997).

44. Cheng, Y., Grigorieff, N., Penczek, P. A. & Walz, T. A primer to single-particle cryo-electron microscopy. Cell 161, 438–449, doi:10.1016/j.cell.2015.03.050 (2015).

45. Chen S. W. et al. Structural characterization of toxic oligomers that are kinetically trapped during alpha-synuclein fibril formation. Proceedings of the National Academy of Sciences of the United States of America 112, E1994–E2003, doi:10.1073/pnas.1421204112 (2015).

46. Zubcevic L. et al. Cryo-electron microscopy structure of the TRPV2 ion channel. Nat Struct Mol Biol 23, 180-+, doi:10.1038/nsmb.3159 (2016).

47. Hite R. K. et al. Cryo-electron microscopy structure of the Slo2.2 Na+-activated K+ channel. Nature 527, 198-+, doi:10.1038/nature14958 (2015).

48. Lendel C. et al. A hexameric peptide barrel as building block of amyloid-beta protofibrils. Angew Chem Int Ed Engl 53, 12756–12760, doi:10.1002/anie.201406357 (2014).

49. Rosenman, D. J., Clemente, N., Ali, M., Garcia, A. E. & Wang, C. Y. High pressure NMR reveals conformational perturbations by disease-causing mutations in amyloid beta-peptide. Chem Commun 54, 4609–4612, doi:10.1039/c8cc01674g (2018).

50. Foroutanpay B. V. et al. The Effects of N-terminal Mutations on beta-amyloid Peptide Aggregation and Toxicity. Neuroscience 379, 177–188, doi:10.1016/j.neuroscience.2018.03.014 (2018).

51. Zirah S. et al. Structural changes of region 1-16 of the Alzheimer disease amyloid beta-peptide upon zinc binding and in vitro aging. Journal of Biological Chemistry 281, 2151–2161, doi:10.1074/jbc.M504454200 (2006).

52. Kawahara, M., Arispe, N., Kuroda, Y. & Rojas, E. Alzheimer’s disease amyloid beta-protein forms Zn2+-sensitive, cation-selective channels across excised membrane patches from hypothalamic neurons. Biophysical Journal 73, 67–75, doi:Doi 10.1016/S0006-3495(97)78048-2 (1997).

53. Tekirian T. L. Commentary: A beta N-Terminal Isoforms: Critical contributors in the course of AD pathophysiology. J Alzheimers Dis 3, 241–248 (2001).

54. Lee J. et al. Amyloid beta Ion Channels in a Membrane Comprising Brain Total Lipid Extracts. Acs Chemical Neuroscience 8, 1348–1357, doi:10.1021/acschemneuro.7b00006 (2017).

55. Diaz, J. C., Simakova, O., Jacobson, K. A., Arispe, N. & Pollard, H. B. Small molecule blockers of the Alzheimer Abeta calcium channel potently protect neurons from Abeta cytotoxicity. Proc Natl Acad Sci U S A 106, 3348–3353, doi:10.1073/pnas.0813355106 (2009).

56. Arispe, N., Diaz, J., Durell, S. R., Shafrir, Y. & Guy, H. R. Polyhistidine Peptide Inhibitor of the Aβ Calcium Channel Potently Blocks the Aβ-Induced Calcium Response in Cells. Theoretical Modeling Suggests a Cooperative Binding Process. Biochemistry 49, 7847–7853, doi:10.1021/bi1006833 (2010).

57. Capone R. et al. All-D-Enantiomer of beta-Amyloid Peptide Forms Ion Channels in Lipid Bilayers. J Chem Theory Comput 8, 1143–1152, doi:10.1021/ct200885r (2012).

58. Fantini J. & Yahi N. Molecular insights into amyloid regulation by membrane cholesterol and sphingolipids: common mechanisms in neurodegenerative diseases. Expert Rev Mol Med 12, doi:ARTN e2710.1017/S1462399410001602(2010).

