## Supplementt Fig. S1 and S2 for "Theory of concentric β-barrel structures: models of amyloid beta 42 oligomers, annular protofibrils, and transmembrane channels"

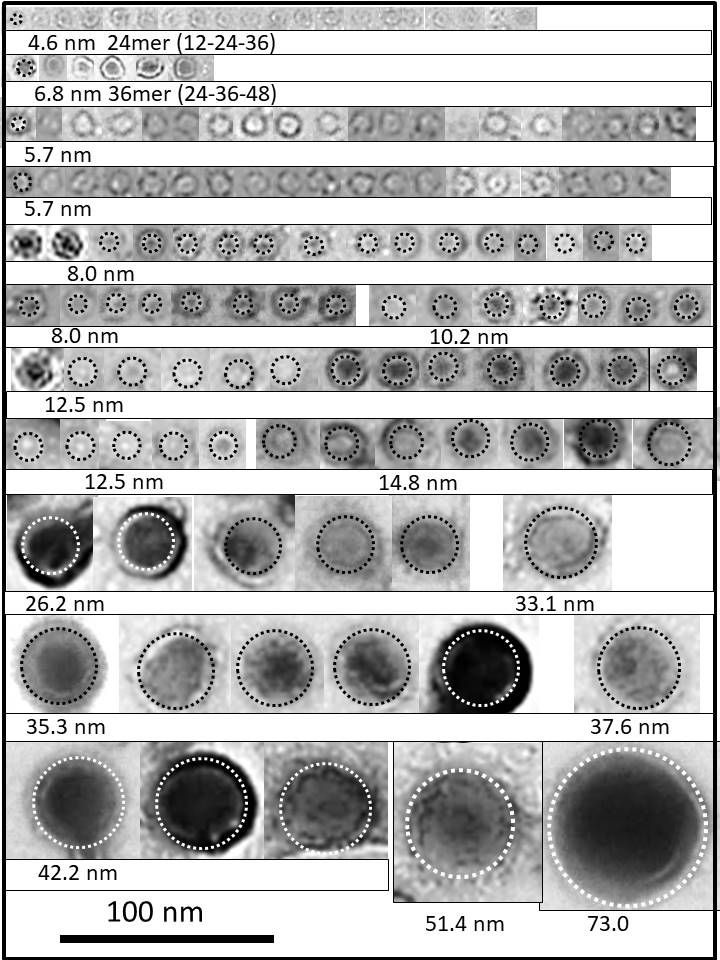

Supplement Figure S1 Additional images of sAPFs not shown in Figure 14. The number of monomers postulated to comprise each category of sAPF is listed below each row.

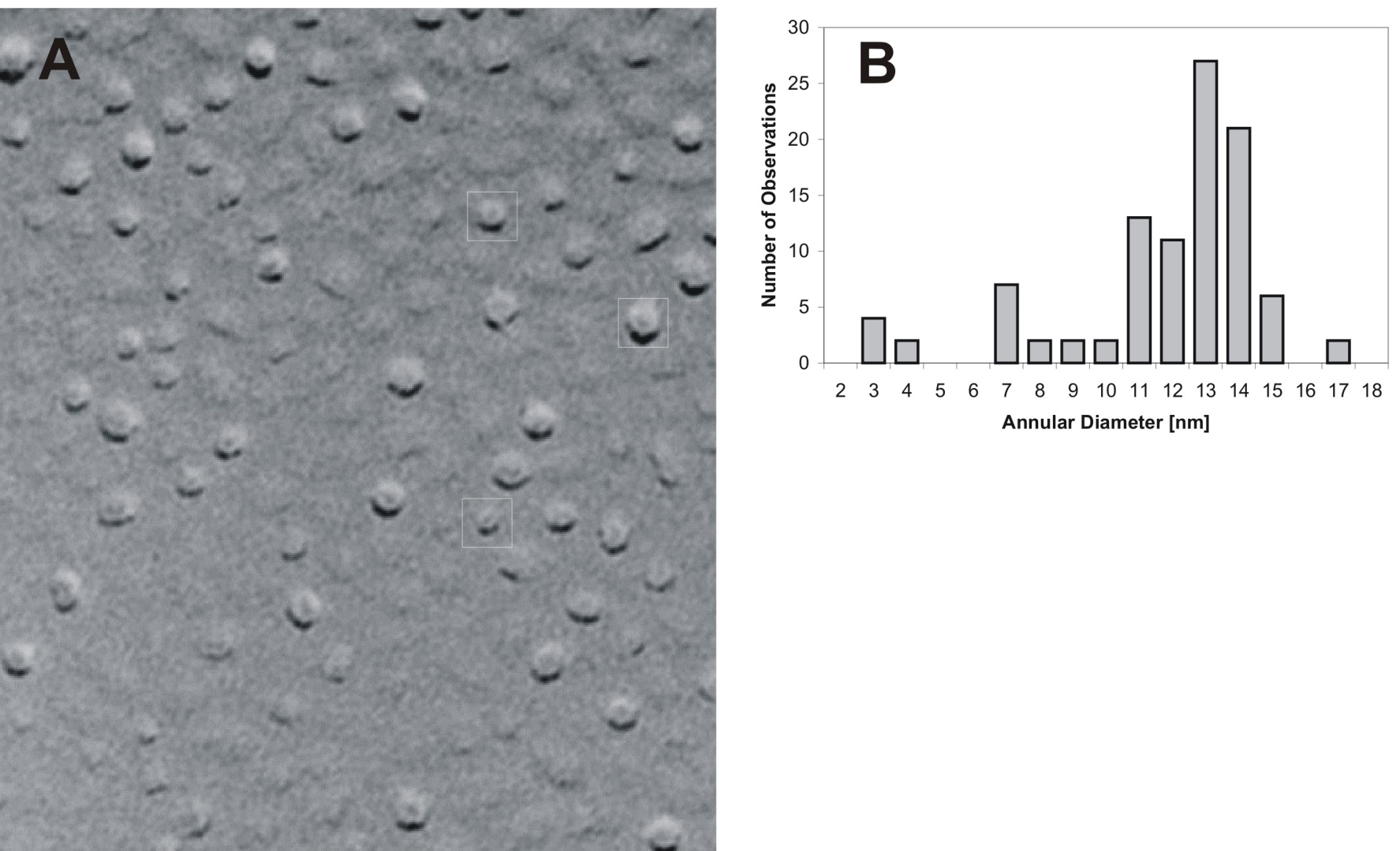

Figure S2. The freeze fracture micrograph of transmembrane Aβ42 assemblies provided by Nelson Arispe.
